# Evolutionary inference across eukaryotes identifies multiple pressures favoring mitochondrial gene retention

**DOI:** 10.1101/037960

**Authors:** Iain G. Johnston, Ben P. Williams

## Abstract

Since their endosymbiotic origin, mitochondria have lost most of their genes. Although many selective mechanisms underlying the evolution of mitochondrial genomes have been proposed, a data-driven exploration of these hypotheses is lacking, and a quantitatively supported consensus remains absent. We developed HyperTraPS, a methodology coupling stochastic modelling with Bayesian inference, to identify the ordering of evolutionary events and suggest their causes. Using 2015 complete mitochondrial genomes, we inferred evolutionary trajectories of mitochondrial DNA (mtDNA) gene loss across the eukaryotic tree of life. We find that proteins comprising the structural cores of the electron transport chain are preferentially encoded within mitochondrial genomes across eukaryotes. A combination of high GC content and high protein hydrophobicity is required to explain patterns of mtDNA gene retention; a model that accounts for these selective pressures can also predict the success of artificial gene transfer experiments in vivo. This work provides a general method for data-driven inference of the ordering of evolutionary and progressive events, here identifying the distinct features shaping mitochondrial genomes of present day species.

## Introduction

Mitochondria are the result of an endosymbiotic event [1], where a free-living organism resembling an *α*-proteobacterium was engulfed by another cell billions of years ago. Since this event, a pronounced loss of mitochondrially-encoded genes has occurred, with genes either transferred to the host nucleus or lost completely [2, 3, 4, 5]. This endosymbiosis and the subsequent evolution of mitochondrial genomes are among the most important processes in biological history, giving rise to eukaryotic life and hypothesised as facilitating the higher energy output required for the evolution of complexity and multicellularity [6].

The precursor mitochondrion is estimated to have possessed thousands of genes [7]. Present-day mitochondrial genomes retain only a small and highly variable number of these: malarial parasites possess only three protein-coding genes in their mtDNA [8] and some protists possess over sixty [9] (human mtDNA encodes thirteen protein-coding genes [10]). Some eukaryotes completely lack mtDNA, representing the limiting case in our picture of mitochondrial gene loss; there is intriguing evidence that changes in nuclear DNA can compensate for a lack of mtDNA in some of these cases [11]. Despite the importance of mitochondrial evolution in diverse fields including phylogenetics [12] and human medicine [10], the forces influencing mtDNA content are still debated and poorly understood, limiting our understanding of the evolutionary history of eukaryotic bioenergetics. Dual questions exist, both as to why so many genes have been lost from the mitochondrion, and why some genes are retained in mtDNA [13].

The first question, why genes are lost from mitochondrial genomes, has a set of answers that are largely agreed upon [14], and several plausible mechanisms [15, 16, 17]. Possible selective advantages conferred by the nuclear encoding of organellar genes include the avoidance of Muller’s ratchet (the irreversible buildup of deleterious mutations) [12, 13], protection from mitochondrial mutagens [18], and enhanced fixing of beneficial mutations [14, 13].

Answers to the second question, why some genes are retained in organelles, remain more elusive. The simplest possibility, which is sometimes assumed, is the ‘null hypothesis’ that gene loss is uniform and random, with no particular link between gene properties and retention in mtDNA. Many possible alternative hypotheses are currently discussed, with two possibilities in particular finding the most qualitative support [19]. The first proposes that highly hydrophobic proteins, if produced remotely, are difficult to import and sort across membranes [14, 20], or readily mistargeted to other organelles [21, 22], favouring the organellar retention of genes encoding these hydrophobic proteins, though the hypothesis is debated [23]. The second hypothesis is known as ‘colocalisation for redox regulation’ (CoRR [23]), suggesting that the retention of key organellar genes allows beneficial localised control of energetic machinery, so that the performance of individual organelles can be optimised with out affecting the whole-cell population. Recent work has supported this hypothesis, showing that genes encoding subunits central to the assembly of the mitochondrial ri-bosome are most retained [24]. Other hypotheses include the possible toxicity of some gene products in the cytosol [25], and differences between the genetic code of the mitochondrion and the nucleus leading to difficulties in interpretation [14]. Additional constraints on mitochondrial gene evolution have been identified including patterns in GC skew, suggested to result from asymmetric mutation pressure [26, 12], GC content, suggested to influence the free energy and thus stability and mutational susceptibility of mtDNA [27], and gene expression, suggested to modulate selective pressure on individual gene sequences in animals [28].

Quantitative evidence supporting one or more of these hypotheses over others is currently absent, with qualitative debate common in the field. Existing studies have analysed specific gene loss events [29], proposed bioinformatic approaches for the analysis of sequence-level features in organellar genomes [30, 31], and explored purifying selection at the mtDNA nucleotide level in animals [28], and, in a broad-ranging study, explored the phylogenetic history and structure of mtDNA using gene clusters and intron structure [32]. However, we are unaware of any existing quantitative approach that identifies the evolutionary pressures behind the highly variable patterns of presence and absence of mtDNA genes across the whole of the eukaryotic tree of life. Although important progress has been made at a sequence level (Ref. [33] takes a phylogenetic approach in studying the sequence evolution of mitochondrial ribosomes across a diverse range of taxa), molecular phylogenies are not necessarily the optimal tools to answer broader questions about the evolution of genome structure [34], particularly given the broad range of taxa and possible parallel evolutionary pathways involved [35, 24]. To make progress with this complex evolutionary system, we developed highly generalisable stochastic and statistical machinery for inference of evolutionary dynamics, and applied it to a large dataset of over 2000 sequenced mtDNA genomes across eukaryotic life, reconstructing the evolutionary history of mitochondrial genomes. We compiled a set of genetic features corresponding to existing and novel hypotheses on the pressures driving mtDNA evolution, and used Bayesian model selection to identify those features, and thus the corresponding pressures, most likely to give rise to the inferred dynamics. We will show that a combination of GC content, hydrophobicity, and energetic centrality (each of which provides independent explanatory power) accounts for most of the variability in observed mtDNA structure and predicts the success of artificial gene transfer experiments.

## Results

### HyperTraPS: an algorithm for sampling rare evolutionary paths on a hypercubic transition network

To explore the evolutionary processes that give rise to present-day patterns of mtDNA genes across taxa, we constructed a general model for mtDNA evolution that allowed us to amalgamate and unify the large volume of genomic data available (see Methods). This model includes a description of every possible pattern of presence and absence of all 65 protein-coding mtDNA genes that we consider. These patterns are represented by binary strings of length *L*, where a 0 as the *i*th character corresponds to the ith gene being absent, and a 1 at that position corresponds to that gene being present. We will use this picture to represent the transitions that make up the evolutionary history of mtDNA (Fig. 1A). We allow evolutionary transitions between states that differ by one trait, so that individual genes are lost one-by-one: however, simultaneous loss of several genes can be captured by this model, and corresponds to an equal probability that each gene is lost at a given timestep. Each transition has a given *intensity*, and the probability of a given transition from some state is proportional to that transition’s intensity (normalised by the sum of intensities of all transitions leading away from that state). Evolution is modelled as a walk from the state of all 1s to the state of all 0s, with individual transitions along the walk occurring randomly, weighted by intensity.

**Figure 1:**
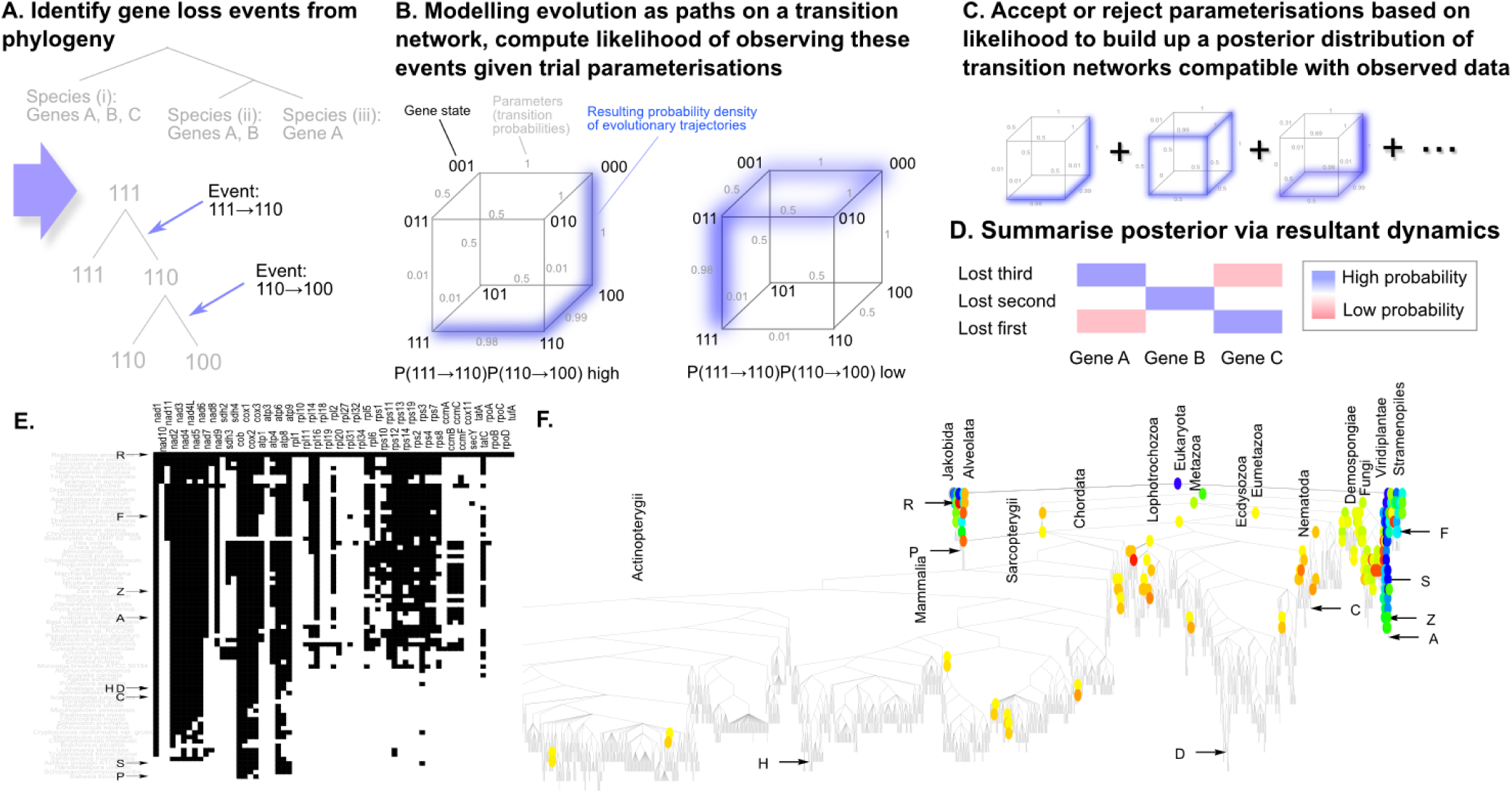
Illustration and source data for HyperTraPS inference of mitochondrial gene loss ordering. (A) Gene loss events are identified by inferring ancestral states on a given phylogeny, providing a set of observed transitions between gene states. (B) An evolutionary space defined by the presence or absence of *L* = 3 traits and parameterised by probabilities of transitions between these states. If our source data reveals two evolutionary transitions 111 → 110 and 110 → 100, the parameterisation on the left is more likely that that on the right, as it supports evolutionary trajectories that are likely to give rise to those observations. HyperTraPS is used to calculate the associated likelihood, determining which parameterisation are accepted (perhaps left) and which are rejected (perhaps right). (C) MCMC is used to build a posterior distribution of parameterisations based on the associated likelihood of observed transitions, producing an ensemble of possible evolutionary landscapes. (D) This posterior distribution is then summarised by recording the probability with which a given gene is lost at a given ordering on an evolutionary pathway. (E) Illustration of the distinct mitochondrial gene sets present in the source dataset, ordered vertically from highest to lowest gene content. Rows are genomes, columns are mitochondrial genes (an example species is given in grey for each genotype). Black and white pixels represent present and absent genes respectively. (F) A taxonomic relationship between organisms used in this study. Each leaf is an organism in which the set of present mitochondrial genes has been characterised. Coloured pairs of nodes denote those ancestor/descendant pairings where a change in mitochondrial gene complement is inferred to have occurred, with hue denoting the number of protein-coding genes present from blue (maximum 65) to red (minimum 3). Single letters in (E, F) denote positions of some well-known organisms by initial letter: *(H) Homo sapiens, (R) Reclinomonas americana, (P) Plasmodium vivax, (F) Fucus vesiculosus, (Z) Zea mays, (S) Saccharomyces cerevisiae, (A) Arabidopsis thaliana, (C) Caenorhabditis elegans, (D) Drosophila melanogaster.*

For example, consider a simple system restricted to *L* = 3 genes. The evolutionary space consists of eight states and ten possible transitions, illustrated in Fig. 1B. If the 111 → 110 transition has intensity 0.98, and the 111 → 101 and 111 → 011 transitions each have intensity 0.01 (Fig. 1Bi), we expect 98% of evolutionary processes to follow the initial step 111 → 110. Subsequent steps from 110 will likewise occur with probability proportional to their relative intensities, as illustrated in Fig. 1B: in Fig. 1 Bi a step from 110 is highly likely to be 110 → 100 due to that transition’s high intensity, whereas in Fig. 1 Bii (with a different set of intensities) a step from 011 is equally likely to be 011 → 010 or 011 → 001, as these transitions have equal intensity. The reader will note that the graphical representations of this model in Fig. 1 B have a cubic structure: as *L* increases this structure expands to become an *L*-dimensional hypercube.

The goal of this section is to construct an algorithm that allows us to compute how likely an evolutionary path within this modelling framework is to visit two given states. For example, consider the case where in our *L* = 3 example above strings describe the presence or absence of *geneA, geneB* and *geneC* in that order. We may have found that an ancestor possessed *geneA, geneB*, and *geneC* (111), and a descendant possesses *A* and *geneB* (110). It is trivial in this case to see that, for the parameterisation in Fig. 1 Bi, the probability of a given evolutionary process visiting state 111 and then state 110 is 0.98. But when L is higher and there are many different paths through which a given state may be visited (or avoided), it becomes prohibitively difficult to compute the probability of every appropriate path. Our approach provides an efficient way to estimate the probability of observing a given pair of states, given a particular parameter set of intensities. Briefly, it accomplishes this by sampling the set of paths that start at the first state and finish at the second, but preferentially sampling the paths which are most likely. This preferential sampling is accomplished firstly by ensuring that no sampled paths involve transitions to states from which the second desired state cannot be reached, and secondly by choosing each step on a sampled path proportionally to its intensity. The amount of bias introduced by this preferential sampling is recorded at each step in the sampled path, and is corrected for to yield the probability required.

In more detail, we work with the ‘evolutionary space’ described in Methods, consisting of a set S of states comprising all possible patterns of presence and absence of L traits under consideration, and a set of transition probabilities *π_s→t_* between each pair of states *s* and *t*, determining the stochastic dynamics of evolution (Fig. 1 A). To compute the probability of observing given evolutionary transitions, we require an estimate of the probability *P*(*s* → *t*|π) that an evolutionary trajectory on a network π passes through a ‘target’ state *t*, given that it passed through a ‘source’ point *s*. This probability is generally hard to compute for a given transition network due to the large number of possible paths that lead from *s* to *t*, since they may differ in any of *L* positions.

We show in the Supplementary Information that an estimate for *P*(*s* → *t*) can be efficiently computed. Consider a path *c*, consisting of a set of states beginning at *s* and ending at *t*, constructed as follows. In the ith path step *c^i^*, identify the set *T(c^i^)* of all accessible states that are compatible with *t* (meaning that they do not lack genes that *t* possesses). Choose the next state *c*^*i*+1^ from *T*(*c*^i^) according to the associated transition probabilities normalised over *T*(*c*^i^) alone. The advantage of this sampling approach is that we can simulate trajectories guaranteed to start at *s* and end at *t*, thus avoiding wasting computational time on trajectories that do not contribute to the overall sum. We underline that our approach does not introduce bias between these pathways – it solely serves to focus computational energy on appropriate pathways. An estimate of *P*(*s* → *t*) is then efficiently given by averaging the quantity Π_*i*_ *P* (*c^i^* → o ∊ *T* (*c^i^*)) over a set of samples, where *c^i^* is the ith state in path *c*, o ∊ *T*(*c^i^*) denotes any member of the set of states accessible from *c^i^* that are compatible with final target *t*, and the sampling is taken over a set of paths, constructed according to the sampling scheme above, starting at *s* and ending at *t* (Fig. S1C). This estimate can be computed using the algorithm below, which to our knowledge has not been previously published; if this is true, we propose the name ‘HyperTraPS’, both standing for hypercubic transition path sampling and referring to the act of forcing trajectories towards specific points on a hypercube. We note that HyperTraPS is in a sense more comparable to the Transition Interface Sampling or Forward Flux Sampling approaches found in statistical physics than Transition Path Sampling as understood in that context [36].

**Algorithm 1**. *Hypercubic transition path sampling (HyperTraPS)*

1. Initialise a set of *N_h_* trajectories at *s*.
2. For each trajectory *i* in the set of *N_h_*:

a. Compute the probability of making a move to a *t*-compatible next step (for the first step, all trajectories are at the same point and the probability for each is thus the same); record this probability as 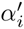.
b. If current state is *s*, set 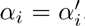, otherwise set 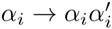
c. Select one of the available *t*-compatible steps according to their relative weight. Update trajectory *i* by making this move.
3. If current state is everywhere *t* go to 5, otherwise go to 2.
4. 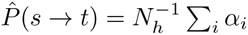

*N_h_*, the number of sampled trajectories, is a parameter of the algorithm. Lower numbers will be computationally cheaper but will give a poorer sampling of possible trajectories and thus a less accurate estimate 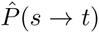. In the Supplementary Information we give a more detailed description of HyperTraPS and show a range of validation checks for its functionality (Fig. S2A-D, S2E). To infer patterns of evolutionary dynamics with HyperTraPS, first a phylogeny is used to identify the evolutionary transitions that have occurred throughout history (Fig. 1 A and Supplementary Information). Then HyperTraPS is used to compute the joint probability of observing these transitions given a trial parameterisation π (Fig. 1 B). An MCMC approach is used to sample parameterisations which are associated with high likelihoods (Fig. 1 C), forming a posterior distribution over π which can be summarised and visualised (Fig. 1 D).

After addressing the specific question of mtDNA evolution, in the Supplementery Information we compare HyperTraPS qualitatively and quantitatively with existing approaches addressing the broad question of characterising the evolution of traits over a phylogeny.

### The inferred pattern of mitochondrial gene loss through evolution

We analyzed the complete annotated mitochondrial genomes of 2015 species from GOBASE [37], identifying the presence and absence of each *R. americana* mito-chondrial gene for each genome. Across the 2015 species, 74 distinct combinations of genes were present (Fig. 1 D). Visible vertical clusters of retained genes include subunits of Complexes I (*nad[X]*), III (*cox[X]*) and V (*atp[X]*), and the small subunit of the mitochondrial ri-bosome (*rps[X]*). We then mapped the differences between each genome onto a phylogeny of all the species in our dataset (Fig. 1 E). We employ two assumptions about mtDNA evolution: first, that mitochondrial gene loss is rare, and second, that mitochondrial gene gain is negligible. These assumptions allow us to reconstruct the mitochondrial genomes at ancestral nodes in the phy-logeny (Supplementary Information; Fig. S1 A-B). To ensure that our approach was robust to potential errors in annotation and phylogeny construction, we repeatedly perturbed our source data and confirmed that our results were comparable across perturbed datasets (Fig. S2E; Methods and Supplementary Information).

We used the HyperTraPS algorithm within a Bayesian MCMC framework to compute the probabilities of different patterns of mtDNA gene loss, given observed changes in mtDNA across the Tree of Life and uninformative uniform priors on transition probabilities. Fig. 2 shows a summary of these results, illustrating the probability with which a given gene is lost at a given step in time from a ‘full’ genome containing all genes found in *R. americana*, to an ‘empty’ mitochondrial genome containing no genes. The figure heuristically represents possible pathways for the evolutionary history of a ‘typical’ mitochondrial genome in an evolving eukaryotic lineage. The pattern of mitochondrial gene loss is remarkably structured and non-random, rejecting the possibility that the genes retained are shared across many species by chance. The inferred structure also quantitatively supports intuitive observations, for example, cytochrome b (*cob*) is observed in almost every known mitochondrial genome, whereas the secY-independent protein translo-case component *tatA* is only observed in *R. americana.* We broadly observe three classes of genes: early loss (including Complex II *sdh[X]* genes, many ribosomal *rps[X]* and *rpl[X]* genes); intermediates present in a variety of taxa (including plants and fungi) but lost in animals (including more subunits of the mitochondrial ribosome and some Complex V *atp[X]* genes); and highly retained genes (including Complex I *nad[X]* genes, Complex IV *cox1-3* genes and cytochrome b *cob*). Different lineages have clearly experienced different specific gene loss trajectories (for example, *Schizosaccharomyces pombe* poss-eses *cox2* but not *cox3*, and *Babesia bovis* possesses *cox3* but not *cox2*, Fig. 1 B), but the probabilistic trend observed in Fig. 2 holds broadly across eukaryotes.

**Figure 2:**
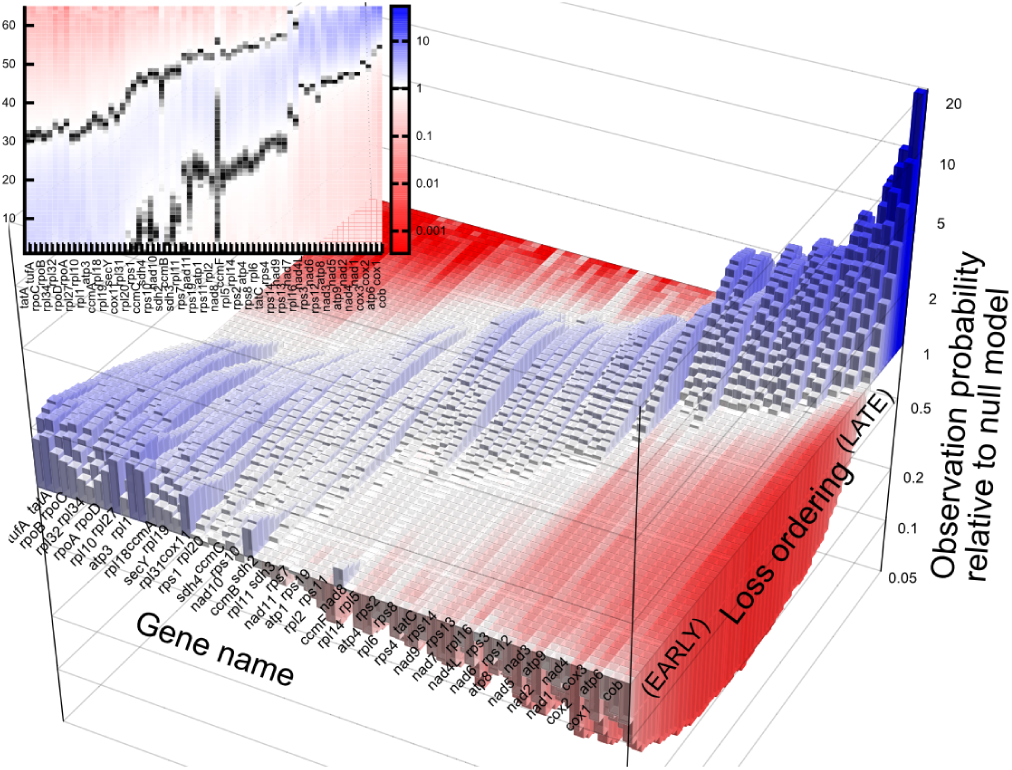
The inferred ordering of mitochondrial gene loss is highly structured and non-uniform. The probability that a given gene is lost at a given time ordering in the process of mitochondrial gene loss. The at surface in the main plot and black contour in the inset give the probability (*l/L*) associated with a null model where all genes are equally likely to be lost at all times. Blue corresponds to a probability above that expected from this null model; red corresponds to a probability below this null model. Genes are ordered by mean inferred loss time.

### Gene loss in electron transport chain complex proteins

Our inferred order of gene loss shows that genes encoding distinct mitochondrial protein complexes were lost in different ‘phases’ throughout evolutionary time. As a first step in more detailed characterisation of this loss patterning, we sought to examine the order of gene loss within mitochondrial protein complexes to address a long-standing hypothesis for why some genes are retained in organelles. As described in the introduction, the CoRR hypothesis suggests that components of central importance to organelle function are preferentially encoded in the organelle genome, to allow localized control of the assembly of protein complexes [23]. This allows individual mitochondria to adjust the stoichiometry of ETC complexes in response to demands or stresses, without affecting the regulation of other mitochondria within the same cell. We reasoned that protein subunits occupying energetically central positions within their complexes likely exert the most control over complex assembly [38] and thus provide a means to test this hypothesis.

We analyzed the crystal structures that have been determined for ETC complexes II, III and IV (see Methods). We found that the genes that encode the proteins with the strongest binding energy (computed using PDBePisa; see Methods) within each complex have invariably been retained in mitochondrial genomes more commonly than those encoding other subunits of the complex (Fig. 3). This relationship is apparent across the entire eukaryotic Tree of Life. The two genes most retained by mitochondrial genomes throughout life, *cob* and *cox1*, are the most energetically central components of their complexes (Complex III and Complex IV respectively). The *sdh[2-4]* genes controlling Complex II, and *cox[2-3]*, which play an important role in Complex IV but are less central than *cox1*, represent intermediate points on this spectrum. Many organisms do not encode any Complex II subunits in mtDNA: those that do so retain the energetically most central *sdh[2-4]* genes and not *sdh1. cox[2-3]* are retained in the mitochondria of most but not all organisms. By contrast, the energetically more peripheral gene products are invariably encoded by the nucleus. This link between biophysical and evolutionary features of mitochondrial genes is strongly consistent with the CoRR hypothesis, supporting its applicability to organellar genomes of multiple and diverse taxa across the tree of life.

**Figure 3:**
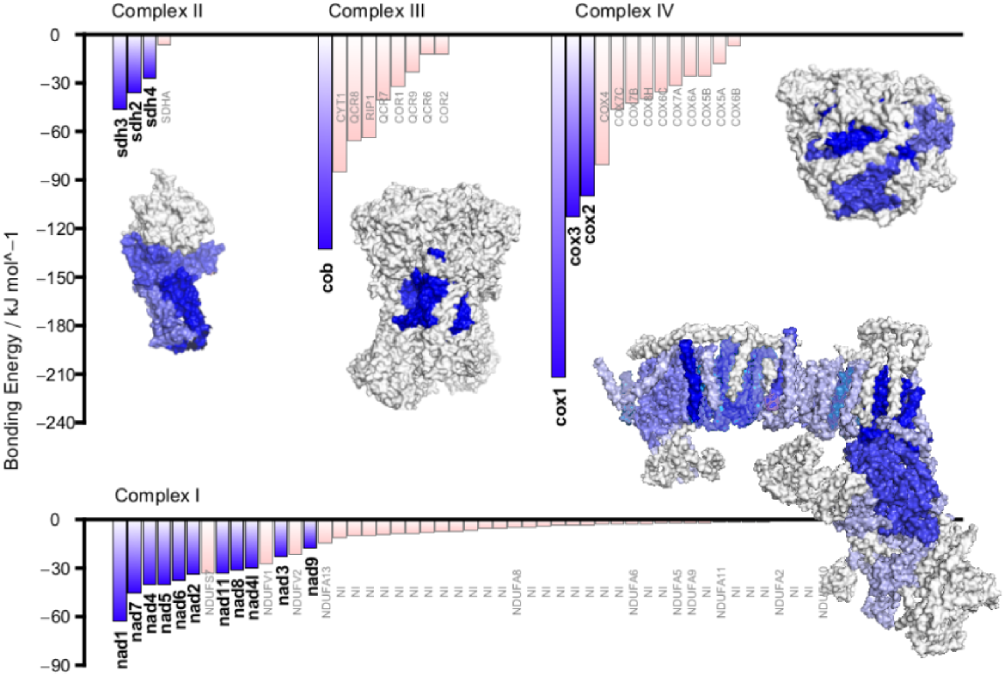
Assembly centrality, gene retention, and colocalisation of redox regulation. Interaction energy of subunits within Complexes I-IV computed with PDBePISA (see Methods). Blue bars denote subunits encoded by mtDNA in at least one eukaryotic species; pink bars denote subunits always encoded in the nucleus. Inset crystal structures show the location of mtDNA-encoded subunits in blue (darker shades show higher interaction energies).

During the preparation of this report, crystal structure data for ETC complex I became available on the PDB [39], affording a valuable opportunity to independently test the prediction of our model. We computed interaction energies for Complex I subunits and found that protein products encoded by mtDNA genes also display the strongest interaction energies, consistent with the pattern we observed with other complexes (Fig. 3). As is the case with the other ETC complexes, the Complex I subunit with the strongest interaction energy, *nad1*, is one of the most commonly retained genes in mitochon-drial genomes across all taxa.

### GC content and protein hydrophobicity are both required to predict mitochon-drial gene retention

We next sought to examine additional factors that predict the probability that a given gene is retained in the mitochondrial genome. To do this, we gathered data for ten properties of mitochondrial genes (or their encoded proteins) that have been hypothesized to influence the probability that a given gene is retained in the mitochon-drial genome (Supplementary Table I). These hypotheses can be viewed as predicting a strong link between the corresponding gene property and the propensity with which a gene is lost from mtDNA. Our characterisation of the probability with which a gene is lost in a given ordering allows us to quantify the strength of these hypothesized links within a probabilistic framework.

We used a linear model approach to explore the ability of features of mitochondrial genes, or the proteins they encode, to predict the inferred patterns of gene loss shown in Fig. 2. As described in Methods, we performed our Bayesian model selection approach with two datasets, one corresponding to genetic features in *R. americana* alone and one with features averaged across taxa. This approach allows us to control both for specific properties of *R. americana* and for cross-species variation. As shown in this section and the Supplementary Information, results from both datasets are very comparable, illustrating the robustness of our findings.

The resultant posterior probabilities over model structure displayed a striking favouring of GC content and protein product hydrophobicity as predictors of whether or not genes are retained in mitochondrial genomes; GC content (B) and hydrophobicity (C) appear in 97-98% of inferred model structures using both *R. americana* gene properties (Fig. 4A) and averaged gene properties (Supplementary Information; Fig. S3A-C). Few other properties are so represented, with the exception of strandedness (J), which appears in 56% of inferred models using averaged gene properties (but few models using *R. americana* properties). Results involving uniform priors on model structure, and using an expanded set of features, were compatible with these findings (Fig. S3A-C; Supplementary Information). The absence of other features in favoured models illustrates a lack of support for other retention determinants including, for example, features related to genetic code variability and mutational robustness. We did not find strong evidence that gene expression is directly related to mtDNA gene retention (Supplementary Information; Fig. S3D), although this analysis was based on a smaller dataset limited by available expression data (see Methods and Supplementary Information).

**Figure 4:**
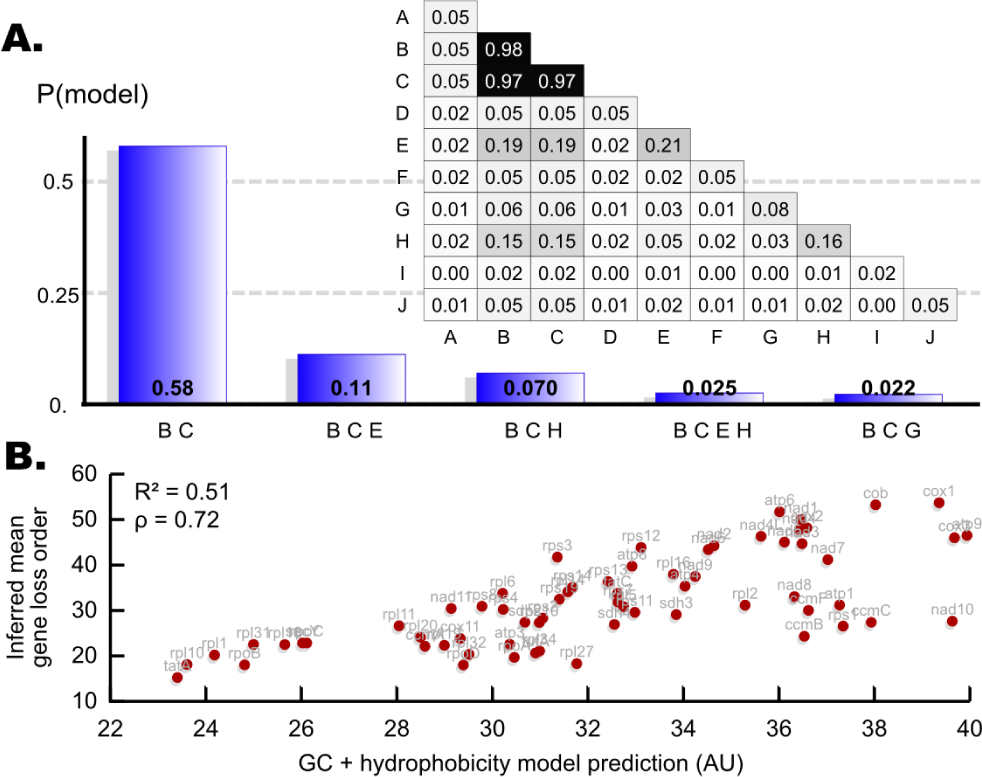
Features predicting mitochondrial gene retention. (A) Bars show posterior probabilities for individual models for mitochondrial gene loss based on gene properties in *R. americana*, given the inferred gene loss patterns and a prior favouring parsimonious models. Inset matrix show the posterior probability with which features are present in these models for gene loss (diagonal elements correspond to a single feature, off-diagonal elements to a pair of features). Labels A-J denote model features as described in Methods; B is GC content and C is protein product hydrophobicity. (B) Predicted loss ordering from GC & hydrophobicity model (horizontal axis) against inferred mean loss ordering (vertical axis).

Our analysis provides strong support for genes with high GC content *relative to other genes in the same organism* being preferentially retained in mtDNA. As we discuss in detail in the Supplementary Information, patterns of GC content vary dramatically between species (Fig. S4; [27]), but this *inter*-species variability is never incompatible with the *intra*-species favouring of GC-rich genes. Notably, at a sequence level, we have found that protein-coding genes in mtDNA can in some species display a bias *against* GC content, in the sense that GC-poor codons are used more often than GC-rich codons to encode a given amino acid (Supplementary Information; Fig. S4). This is most likely due to the strong asymmetric mutational pressure arising from the hydrolytic deamination of cytosine into uracil, which is converted into thymine. This mutational pressure likely enriches GC-poor codons in organellar genomes [26]. Our results suggest a tension between this ‘entropic’ mutational drive at the sequence level (decreasing genome-wide GC content) and a selective drive at the genomic level (retaining genes with higher GC content).

To further elucidate the nature of this tension, we examined the GC content of individual genes at non-synonymous and synonymous positions in *R. americana* and averaged across taxa. We found that GC content was lower at synonymous positions than at nonsynonymous positions in *R. americana*, but that this difference largely disappeared in the taxa-averaged data (Fig. S4B). This difference in *R. americana* is consonant with a link between GC content and structural conservation: nonsynonymous sites retain their GC content as conservation pressure balances the asymmetric mutation pressure, while synonymous sites are free to decrease GC content in response to asymmetric mutation. In this picture, the link between GC content and gene retention could arise indirectly through another relationship: genes most retained in mtDNA being most highly conserved (perhaps due to their structural importance in protein complex assembly, as above). However, the absence of this signal in the taxa-averaged data suggests that this relationship may not hold across all the species we consider. We also tested the link with conservation by using nonsynonymous, synonymous, and total GC content as different terms in our model selection process (Supplementary Information, Fig. S4C). The model selection process always favoured total GC content, suggesting that the conservation link, while explanatory in *R. americana*, does not represent the only way in which GC content is linked to gene retention (see Discussion).

The parameterisation of the most common model always favoured the retention of genes encoding proteins with high hydrophobicity and high GC content. In Fig. 4 B we illustrate a specific parameterisation of this model, chosen to optimise the least-squares difference between model prediction and mean inferred ordering (from Fig. 2). This parameterisation shows a strong gene-by-gene correlation with the mean inferred loss ordering, with Pearson’s *R*^2^ = 0.51 (*p* ≃ 2 x 10^−11^, see Supplementary Information for interpretation) and Spearman’s *ρ* = 0.72. The model also strongly correlated with observation frequencies of mtDNA genes in the original dataset (Fig. 1 B, S3G).

Intuitively, it may be expected that the signals associated with GC content, hydrophobicity, and energetic centrality may reflect different aspects of the same fundamental feature: GC-rich codons encode hydrophobic amino acids, and hydrophobic proteins are likely to occupy ‘core’ positions in complexes and within membranes. However, there is evidence that these features are at least somewhat decoupled. Fig. S3F shows an absence of strong correlations between the three features for the genes we consider: Complexes II and IV display a moderate correlation between scaled energetic centrality and GC content (though sample size is limited by the small number of subunits), but no relationship exists for other complexes or hydrophobicity. Additionally, our model selection process will discard redundant information; if all the predictive power associated with GC content was already present in the hydrophobicity data, GC content would not be identified as a joint factor in the selected models. While the links above are certainly true in some contexts, we cannot avoid the conclusion that independent features associated with GC content, hydrophobicity, and energetic centrality all contribute to gene loss propensity. A combination of these, as opposed to any single feature, is therefore required to account for observed patterns of mtDNA gene retention.

Our findings give rise to a range of predictions on cellular and evolutionary scales. At a systematic level, observation of an organism with mtDNA encoding GC-poor, low-hydrophobicity, energetically peripheral genes and not GC-rich, hydrophobic, energetically central genes would stand against our theory. On a cellular level, our theory predicts that attempts to encode GC-rich, hydrophobic, energetically central genes in the nucleus rather than in mtDNA will adversely affect cell functionality. This prediction can be tested through artificial gene transfer experiments, and indeed, several recent and intriguing experimental studies in *S. cerevisiae* attempting to artificially transfer mtDNA genes to the nucleus allow us to test our theory. To our knowledge, such experiments have been attempted for genes *cob* [40] (unsuccessful), *atp9* [41] (unsuccessful but introduction of a nuclear-encoded version from another species succeeded), *cox2* [42] (successful after a small structural modification), *atp8* [43] (successful), and *rps3* (successful) [44]. This relative ordering of dificulty matches the predictions made from our theoretical treatment as observed in Fig. 5, supporting our GC/hydrophobicity theory.

**Figure 5:**
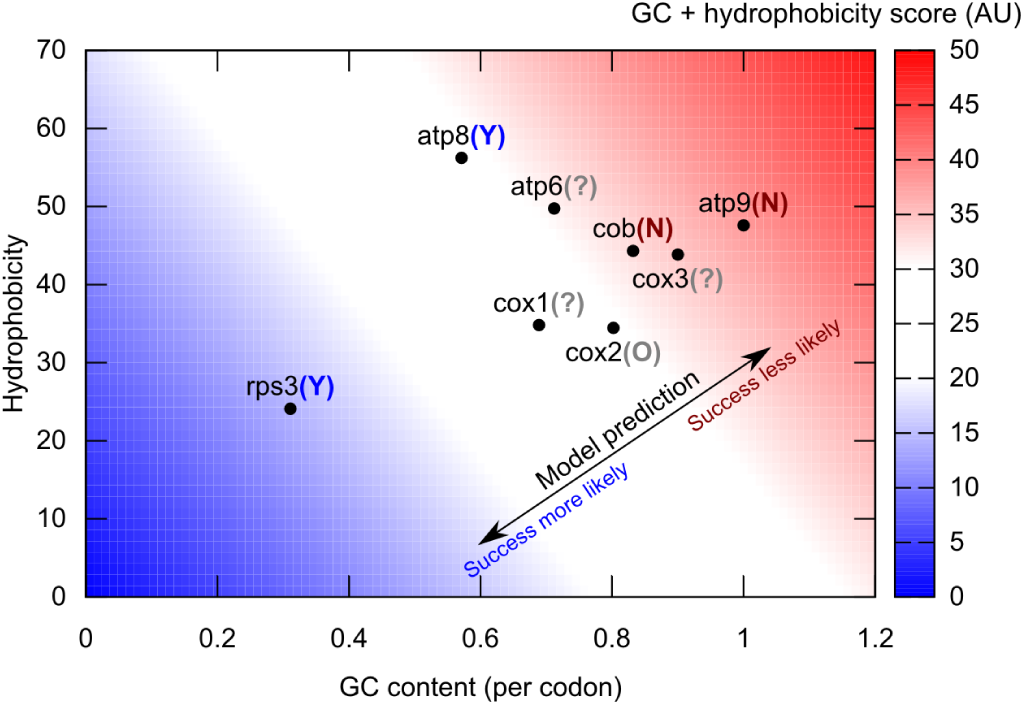
Predicted and observed feasibility of experimental mito-nuclear gene transfer in S. cerevisiae. The protein-coding mtDNA genes in *S. cerevisiae*, plotted on a space of GC content and protein hydrophobicity. Heat map gives the value of the fitted model from Fig. 4 B. Parentheses give the experimental status of mito-nuclear transfer for the given gene: Y {successful, N {unsuccessful; O {partial success (after structural modification), ? {currently unattempted. The success of transfers follows the model prediction for loss propensity, and predictions for the ease of unattempted gene transfers can be formulated.

## Discussion

We have developed new mathematical machinery combining stochastic modelling with Bayesian inference to infer structure and variability of evolutionary pathways, and applied it to explore the evolution of mtDNA gene content across eukaryotes. In the Supplementary Information we compare HyperTraPS to related approaches in the phylogenetic and cancer progression literature (reviewed respectively in Refs. [45] and [46]) and and show that it offers increased exibility and computational efffciency above existing methods. We believe that our highly generalizable mathematical approach may be used to answer a broad array of questions in biology and medicine, offering the ability to work with a general search space comprising dozens of features, which are unrestricted and may comprise biologically distinct classes (for example, combining physical and genetic characteristics [47], or unrelated clinical phenotypes), and the ability to make probabilistic predictions with quantified uncertainty about behaviour given a particular state, and unmeasured features of samples (verified in a plant study [47]). Our approach can be employed in cases where discrete traits change irreversibly with time (though we are pursuing the extension to reversible transitions), and we anticipate it being of use in fields (with corresponding traits in parentheses) including cancer progression [48] (chromosomal aberrations); antibiotic resistance in tuberculosis [49] (resistance to given drugs); chloroplast evolution [50] (‘tribes’ of organelle genes); and paleontology (discrete morphological traits).

Applied to the question of mtDNA gene loss, we have identified structure in mitochondrial evolution across eukaryotes, involving parallel losses across lineages and fine-grained lineage variation imposed on a broad, predictable cross-species trend. We have focussed on protein-coding mitochondrial genes, to facilitate an analysis of the potential modulating factors of this gene loss. The mitochondrial genes encoding, for example, rRNA and tRNA are likely subject to different evolutionary pressures and will be the target of future investigation.

We have quantitatively demonstrated that genes central to the assembly of ETC protein complexes are preferentially retained in eukaryotic mtDNA. This observation suggests a picture of a controllable subset of ‘core’ genes retained by the mitochondrion to allow localised bioenergetic modulation, with a cytoplasmic pool of ‘periphery’ subunits ready to assemble around any newly-produced core, congruent with recent findings that genes encoding central subunits of the mitochondrial ribosome are preferentially retained in mtDNA [24]. This idea of a localised and controlling relationship between mtDNA and energetic machinery supports the central aspects of the CoRR hypothesis [23] and the picture of locally addressable ‘quality control’ of mitochondria [51, 52]. A related recent hypothesis [6] proposes that the acquisition and subsequent genetic reduction of mitochondria was a major facilitating step in the evolution of complex life, as localised cellular power stations controlled through a small number of genes dramatically increase a cell’s available energy per gene, allowing genetic exploration without sacrificing energy. Our identification of energetically central ‘control’ genes as preferentially retained in mitochondrial genomes supports CoRR across the range of taxa that we consider.

Energetic centrality in ETC complexes provides a suggestion towards why a particular subset of the thousands of genes possessed by ‘proto-mitochondria’ has been retained in modern eukaryotes. A related question is why, within this subset, certain genes are retained in the mtDNA of more species than others. Considering the full range of mtDNA-encoded genes, we have identified two features – GC content, and the hydrophobicity of an encoded protein product – that strongly predict the propensity of a gene to be retained in mtDNA across taxa. Other proposed features modulating retention, such as genetic code differences, gene expression levels, and GC skew have much lower support from our approach, and a link between mtDNA retention and gene expression levels is evident in some but not all species, and likely secondary to the link with GC content (Supplementary Information). We underline that the explanatory contributions of GC content, hydrophobicity, and energetic centrality to gene retention are at least partly independent: each factor has individual explanatory power, suggesting mechanisms associated with each. As discussed by Allen and Cavalier-Smith following Ref. [53], a combination of features is therefore required to account satisfactorily for gene retention patterns, and our results quantitatively describe the explanatory components of this combination for the first time.

If a coupled picture where these features report the same underlying property – hydrophobic, central, GC-richly-encoded subunits obeying CoRR – is not the whole story, what other mechanisms could underlie the observed links? Regarding hydrophobicity, it has been suggested that import of hydrophobic proteins through membranes into the mitochondrion may be limiting [20, 14], though this picture is debated [23]. Additionally, it has been proposed that hydrophobic proteins are more likely to be directed to the endoplasmic reticulum than to the mitochondria when translated in the cytoplasm [21]. Importantly, a recent study has experimentally verified that mitochondrially-encoded proteins are indeed localized to the endoplasmic reticulum when expressed in HeLa cells [22] – thus providing a mechanism for hydrophobic modulation of mtDNA gene retention without invoking the controversial membrane-traversal picture. Our results are consistent with the plausible picture that emerges – that the sub-cellular targeting of hydrophobic proteins is an important selective constraint that has favored retention of some genes in mitochondrial genomes.

The independent contribution of GC content to mtDNA gene retention is less mechanistically clear. We show in Figs. S4B and S4C that a putative link between structural conservation and gene retention may account for some, but not all, of the explanatory power of GC content. We suggest that the remaining connection between total GC content and mtDNA gene retention may be manifest through the physical and chemical properties of mtDNA and derived nucleic acids, and their survival in the damaging environment of the mitochondrion. Firstly, GC content has been hypothesised to modulate longevity through its influence on the thermodynamic stability of the mtDNA molecule [27]: nucleic acids containing more (stronger) GC bonds may be less prone to spontaneous ‘bubble’ formation and thus more protected from environmental mutagens. Secondly, adenine depletion has sometimes been observed during oxidative stress [54, 55]. MtDNA could thus preferentially utilise GC-rich codons to optimise chemical stability of nucleic acids in the mitochondrion, and to avoid depleting these redox-linked pools under stress, resulting in selection for increased GC content. This hypothesis is experimentally testable, for example by measuring the extent of DNA damage in GC-rich versus GC-poor mitochondrially encoded genes under stress, which we would expect to be exacerbated in mutants for mitochondrial DNA repair pathways. Further, if redox-linked adenine depletion does constitute an important pressure favouring GC content, GC-richer mtDNA haplotypes should experience an advantage under oxidative stress. A confirmatory assay could be performed in several existing models where two different mtDNA haplotypes exist in admixture [56]. In addition to providing the first statistically robust support for long-debated hypotheses regarding mitochondrial gene loss, and identifying the combination of factors governing this process, our study thus also yields simple, experimentally testable predictions from a complex set of evolutionary transitions.

### Experimental Procedures

#### Data mining and curation for mtDNA gene content

The jakobid protozoan *Reclinomonas americana* has a large mitochondrial genome [9] which includes orthologues of every gene found in any organism’s mitochondrial genome (with very few exceptions [57]). We use the set of *L* = 65 identified protein-coding genes in the mtDNA of *R. americana* as our reference set (Supplementary Information). We represent mitochondrial genomes from across all kingdoms of eukaryotic life as strings of *L* presence or absence markers, one for each protein-coding gene in the *R. americana* genome. To obtain these representations, we use the full set of data from GOBASE, the organellar genome database [37], consisting of annotated genome records for 2 015 species after duplicates were removed. We used the NCBI Taxonomy tool [58] to build an estimated tree of taxonomic relationships between all species recorded in the dataset, and manually verified the topology of this tree with comparison to the Encyclopedia of Life [59] and Tree of Life [60] projects. We then identify the changes that mitochondrial genome content has undergone throughout evolutionary history by comparing inferred ancestral mitochondrial properties with descendant properties (Fig. S1A-B; Supplementary Information). To ensure that our results are robust with respect to perturbations in the details of the constructed phylogeny and this inference protocol, we applied sets of random changes to the resultant dataset and verified that our results were consistent across this set of random changes (Supplementary Information).

#### Representing evolutionary space and transition networks

We consider an underlying ‘evolutionary space’ described by a transition network π*S_i_ → S_j_*, giving the probability of a transition from state *S_i_* to *S_j_*, where states correspond to binary strings of length *L* as above. In any state, we consider transitions corresponding to the loss of exactly one mtDNA gene, restricting *π* to a hypercubic structure (Fig. 1 B). We write π using π_*s* → *t*_ = *Z*^-1^ *P*(lose trait i|state *s*)*I*(*s,t,i*), where *I*(*s,t,i*) is an indicator function returning 1 if *t* is equivalent to s after the loss of trait *i*, and 0 otherwise, and *Z* is a normalising factor to ensure ∑_*t*_ π_*s*_ → _*t*_ = 1. This structure allows us to coarse-grain the pa-rameterisation of the problem by writing P(lose trait *i*|state *s*) = exp (*m_i,j_*_1_ + ∑_*j*_*m_i,j_*_+1_(1 – *s_j_*)), where *m* is an *L* X (*L* + 1) matrix *m*, with elements in the first row *m*_*i*,1_ representing the ‘default’ probability of losing trait *i*, and elements in the subsequent rows *m_i,j_*_+1_ describing how this default probability changes in a state where trait *j* has already been lost. This representation, as discussed in Ref. [61] where this philosophy is also employed, allows us to use *O*(*L*^2^) parameters rather than the full *O*(2^*L*^) set of transition probabilities, while retaining the ability to model both independent gene loss propensities and potential contingencies of the loss of one gene on the presence of others. We illustrate its ability to satisfactorily capture evolutionary behaviour in the Supplementary Information.

#### Inferring mtDNA gene loss ordering

We developed an extended and generalised version of phenotype landscape inference [47], using an algorithm we call HyperTraPS (hypercubic transition path sampling; see Results). We use HyperTraPS to compute the probability of observing the set of mitochondrial gene transitions that we infer from genomic data, given the matrix *m* (see above) representing transition probabilities between different mtDNA states. This probability is used in a Bayesian MCMC algorithm (Fig. 1 A-D; see Supplementary Information for quantitative details) to obtain a posterior distribution on *m.* This posterior distribution is summarised as a posterior distribution over mtDNA gene loss orderings (Fig. 2).

#### Bayesian model selection for features determining mtDNA gene loss propensity

To investigate the potential determining factors that govern mitochondrial gene loss ordering, we compiled a list of many physical and genetic features of mitochondrial genes using sequence data [37] and standard chemical data sources [62] (Supplementary Information). Features included (A) gene length, (B) GC content, (C) hydrophobicity of product, (D) molecular weight, (E) *pK_a_*, (F) energetic cost of production [63], (G) codon universality (the proportion of codons whose interpretation varies across eukaryotes [14]), (H) mutational robustness [13], (I) GC skew, and (J) the strand on which the gene was encoded. As described in the Supplementary Information, this set of features allows us to explore existing hypotheses regarding mitochondrial gene retention (including, for example, hydrophobicity and genetic code differences), while also investigating gene properties that have not previously been explicitly considered in the literature (including GC content and an associated role for nucleic acid stability [27]). We also used RNASeq data from two species with ∼ 30 mtDNA genes (*Lolium perenne L.* [64] and *Phoenix dactylifera L.* [65]) to quantify links between gene retention and expression levels. These species have, to our knowledge, the highest number of mtDNA genes of any species in which gene expression has been quantified; we therefore selected them in order to quantify expression levels (and thus explore potential correlations) for the maximum possible number of mtDNA genes. Full details of how each feature is represented, and their links to individual hypotheses, is presented in the Supplementary Information. For A-J, we created two libraries of genetic properties for protein-coding mtDNA genes: one based upon values derived from *R. americana* (to control against interspecies variation of these properties) and one based upon values averaged over a set of species representing the full range of available eukaryotic diversity (to control against features specific to *R. americana*). We used a Bayesian model selection approach to compare the support for linear models involving combinations of these features given the patterns of mitochondrial gene loss inferred by HyperTraPS. We used two different classes of prior: first, a uniform prior probability over all possible models regardless of the number of features of each, and second, a prior probability exponentially decreasing with the number of features involved, to favor sparser and more parsimonious models.

#### Energetic centrality of ETC complex subunits

To explore the CoRR hypothesis, we investigated the interaction energy associated with a subset of mitochondrial genes in their respective protein complexes, as a measure of the energetic centrality that these genes play in the assembly and structure of each complex [38]. To quantify this measure, we used the PDBePISA tool [66, 67] to analyse the energetic interactions of the subunits in solved structures of electron transport chain complexes in the PDB [68] (specifically, Complex II (PDB 2h88, *Gallus gallus*), Complex III (PDB 3cxh, *Saccharomyces cerevisiae*, Complex IV (PDB 1oco, *Bos taurus)*, and recently Complex I (PDB 4uq8, *Bos taurus))*, and identified the subunits corresponding to genes that are present in our reference set, using annotations associated with each PDB entry, and the UniProt resource [69].

## Acknowledgments

The authors thank A. Earl, N. Jones, A. Monti, and E. Røyrvik for valuable discussions, and four anonymous reviewers for highly valued suggestions about the manuscript.

## Supplementary Information

### Mitochondrial gene set acquisition and curation

The GOBASE database was used to obtain a set of records of all recorded mitochondrial genomes. Each record contained a species name, accession ID, and set of mitochondrial genes present. In the vast majority of cases, these sets formed a subset of the genes present in *Reclinomonas americana*, our reference organism. After duplicates were removed, our dataset included the genomes of 2,015 distinct species.

We considered the set of the following 65 identified protein-coding genes in *R. americana*: *atpl, atp3, atp4, atp6, atp8, atp9, ccmA, ccmB, ccmC, ccmF, cob, cox1, cox11, cox2, cox3, nad1, nad10, nad11, nad2, nad3, nad4, nad4L, nad5, nad6, nad7, nad8, nad9, rpl1, rpl10, rpl11, rpl14, rpl16, rpl18, rpl19, rpl2, rpl20, rpl27, rpl31, rpl32, rpl34, rpl5, rpl6, rpoA, rpoB, rpoC, rpoD, rps1, rps10, rps11, rps12, rps13, rps14, rps19, rps2, rps3, rps4, rps7, rps8, sdh2, sdh3, sdh4, secY, tatA, tatC, tufA.*

The NCBI Common Taxonomy Tree tool was used to construct a taxonomic tree for the species for which data was obtained. We then inferred the barcode of each branching point on the tree simply by applying the OR operator bitwise on each branch’s barcodes. This protocol relies on two assumptions:

1. Mitochondrial gene loss is rare, so if two descending lineages of a common ancestor are observed to lack a gene, we assume that their common ancestor lacks it, as opposed to gene loss occurring convergently in both descendant lineages;
2. Mitochondrial gene *gain* is very rare [57], so if an ancestor has two descendants, one with and one without a gene, it is much more likely that that gene has been lost in one descendant than that it has been gained in the other.

Finally, we recorded the subset of edges on the tree where a change occurred between two connected mitochondrial genomes, from ancestor to descendant (Fig. 1A). Our dataset 𝒟 thus consists of *n_D_* pairs of barcodes {*α_k_*,*d_k_*}, respectively ancestor and descendant in pair *k*.

We note that this approach allows us to account for the large sampling bias in recorded mitochondrial genomes. Many more vertebrate mtDNA sequences have been recorded than any other clade: however, mitochondrial gene sets are largely homogeneous across vertebrates. By only using instances where an independent change between parent and daughter has been observed, we ignore this oversampling and assign equal weights to each event of evolutionary change.

## Inference of evolutionary dynamics

### Mathematical background

We consider a state space consisting of all binary strings of length *L*, and a hypercubic transition network π describing the probability of a transition between two states. We assume that π is structured such as to ensure that a trajectory starting at 1^*L*^ terminates after *L* steps at 0^*L*^, with each step involving a change of one locus in the current string from 1 to 0. The set of transition probabilities in π leading from any node are, by definition, constrained to sum to unity. We write *P*(*b*|*a*; π) for the probability that a system initially at *a* will transition to *b*. The ‘origin’ state *O* ≡ 1^*L*^ is where all evolutionary trajectories begin.

We will work in the picture of a hidden Markov model [70]. Here, the process of evolution in a single lineage corresponds to a trajectory across the hypercube, starting at 1^*L*^ and potentially ending at 0^*L*^. Trajectories emit ‘signals’ at random with a characteristic rate. Each signal is simply the current state of the trajectory. There is thus a constant probability of emitting a signal at any state in a sampled trajectory. If some trajectories are more likely than others, and an ensemble of trajectories is simulated, more emitted signals will be expected from states in common trajectories than from states in rare trajectories.

The fundamental quantity of interest throughout is *the probability that randomly emitted signal(s) from a randomly sampled evolutionary trajectory match an observation.* The product of this probability over all independent observations (see below for discussion of independence) constitutes the likelihood of a given transition network given biological data.

### Dynamics and likelihoods for evolutionary lineages

This subsection shows that the likelihood associated with data 𝒟, a set of *n_D_* observed transitions {*a_i_* → *d_i_*}, is

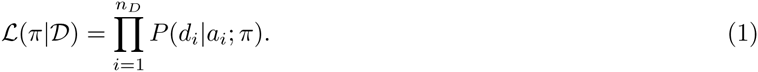

This result may hold intuitively for some readers; they are directed to the next subsection, where an efficient method for computing these probabilities is developed.

In the original application of phenotype landscape inference [47], the properties of observed species were assumed to be the product of convergent evolution, with each patterns of presence or absence the result of an independent evolutionary trajectory. Observations thus consisted of single states, where each observation was independent of all the other observations. The likelihood associated with an observation, as above, is the probability that a randomly emitted signal from a randomly sampled evolutionary trajectory matches the observation. To estimate this probability, random, independent evolutionary trajectories were simulated on π, and the proportion of states in each trajectory that were compatible with an observation was computed. This proportion was then proportional to the probability that a randomly emitted signal from a random trajectory matched the observation, with the constant of proportionality a multiplicative factor accounting for emission probabilities that was independent of model parameterisation and hence vanished in likelihood calculations.

The non-trivial taxonomic relationship between observations in the mitochondrial system means that this picture of independent trajectories leading to independent observations no longer applies. Instead we must consider the pattern of shared histories in the set of observations. For example, the joint probability of two observations known to come from the same mitochondrial lineage cannot be calculated as the joint probabilities of two independently sampled trajectories emitting signals compatible with those observations.

To give a concrete example, consider a system where we have observed the properties of two species: species A with 110 and species B with 100. In the convergent evolution picture, species *A* and species *B* have both evolved independently from a common ancestor O with 111, and the joint probability of these (independent) observations would simply be *P_obs_*(111→110)*P_obs_*(111→100). However, if species B is a descendant of species A, the evolutionary processes are no longer independent: we know that both observations have been made in the same lineage. Hence, the joint probability is *P_obs_*(111→110)*P_obs_*(110→100|111→110). The important difference is that the non-convergent picture is ‘serial’, involving sequential observation probabilities contingent on previous steps, whereas the convergent picture is ‘parallel’, involving independent descents from the original common ancestor.

We have used *P_obs_*(*o* → *o*) for the probability of *making an observation* as opposed to the probability *P*(*o*|*o*; π) of undergoing a transition from one specific state to another. The two are different for two reasons. Firstly, to make an observation of a transition, we require a signal to be emitted at the initial and final state; there is a probability associated with each event, corresponding to the probability of emitting signals from the evolutionary process at a specific time. We write *P_emission_* to denote the probability of emitting signals at the required times. Secondly, the probability of observing a transition is generally proportional to the product of that transition probability and the probability *P_reach_*(*α*; π) of encountering its initial state. This captures the fact that, even if a transition to *b* is certain for a system at *a*, this transition will never be observed if the system never reaches *α*. In general, then, we have

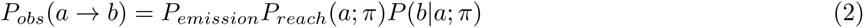

We will ignore the factors of *P_emission_* henceforth because if, as before, signals are emitted randomly and uniformly, independent of model parameterisation and the state of an evolutionary trajectory, *P_emission_* is a constant multiplicative factor that is a function of the data structure alone. This factor will then cancel when likelihood ratios are considered. Emission probabilities are discussed further, with examples, in ‘Signal emission probabilities’ below.

We are left with *P_obs_*(*a* → *b*) = *P*(*b*|*a*;π)*P_reach_*(*a*;π). In the convergent picture, *P_reach_*(*a*;π) vanishes as the initial state is always the origin state *O* where all trajectories start, and *P_reach_*(*O*) = 1.

In the non-convergent picture, we have serial chains of observations, with the ancestral state in each observation being the descendant in another observation, except in the case of the origin state. We thus have a chained product of observation probabilities. For example, the observations *a* → *b*, *b* → *c*, and *c* → *d* in the same lineage give *P_obs_*(*c* → *d*|*b* → *c*)*P_obs_*(*b* → *c*|*a* → *b*)*P_reach_*(*a*; π). Each observation probability is contingent on having already reached the initial state through a different observation, and the initial state dependence thus vanishes for all but the most ancestral transition: for example, the term corresponding to an intermediate observation in the chain above, *P_obs_*(*b* → *c*|*a* → *b*) = *P*(*c*|*b*; π)*P_reach_*(*b*|*a* → *b*) = *P*(*c*|*b*; π) x 1. The origin state is encountered in every trajectory, as in the convergent case, and so the associated term is also unity.

Furthermore, we note that the initial state emission probability in Eqn. 2 is 1 if that state has already been emitted as the final state of a previous transition. Each observation thus only has one emission probability factor (which still cancels as discussed above).

The probabilities associated with evolutionary trajectories in parallel lineages are independent after their common ancestor and can straightforwardly be multiplied. It can then readily be seen that any combination of parallel and serial lineages reduces to a joint observation probability involving only the product of individual transition probabilities (an example is illustrated in Fig. S1A-B). Hence, for our inference purposes, it will suffice to consider the product of probabilities of observed transitions as the likelihood associated with a given dataset. We therefore have, neglecting multiplicative constants:

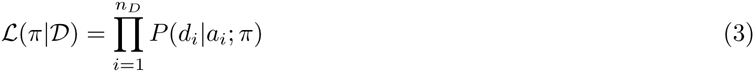

An example of this calculation is given in Fig. S1C.

If all observed transitions *a_i_* → *d_i_* involved single changes, and hence single edges of π, each term in the likelihood function product could straightforwardly be read off from π. However, in general, *a_i_* and *d_i_* may differ at many positions, and many different trajectories may be used to transition between the two. We therefore require a way to calculate the probability of this transition, taking these different possible trajectories into account.

### Efficiently estimating transition probabilities

We consider the problem of computing the probability that an evolutionary trajectory, beginning exactly at source *s*, will lead to the observation of target *t*, given transition matrix π. *Note that we here use t to denote a target state rather than time*. Transition probabilities *P*(*b*|*a*; π) are determined by the edge weights of π; however, for clarity in the working below, we will not write this π dependence explicitly, and will adopt the symbol:

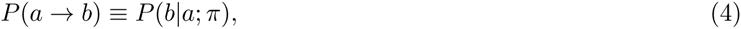

We are thus interested in the probabilities associated with all trajectories that lead from *s* to *t*. Labelling a trajectory consisting of *N* steps as *c* = *c*^0^, *c*^1^, *c^N^*, we have:

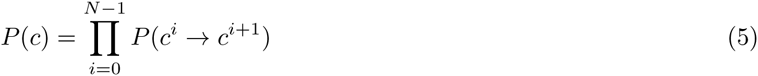

and so

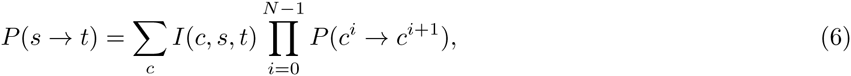

where *I(c, s, t)* is an indicator function returning 1 if trajectory *c* starts at *s* and ends at *t* and 0 otherwise.

Generally we expect this sum to be hard to perform through random sampling, as only a small number of all possible trajectories may pass through *s* and *t*. A very large number of randomly chosen trajectories will then need to be simulated to ensure that we characterise *P*(*s* → *t*).

We instead consider an approach where we constrain the trajectories we simulate to start at *s* and end at *t*, and account for the amount of bias we need to employ to do so.

#### Definition

A state *r* is *t*-compatible if *r_i_* = 1 for all *i* for which *t_i_* = 1.

**Lemma 1**. Each step on a trajectory that will eventually reach *t* must be *t*-compatible, as we cannot reacquire lost traits.

Define *T*(*c^i^*) as the set of *t*-compatible states that can be reached by acquisitions from point *c^i^*. Consider two events: (a) a transition from *c^i^* to *c*^*i*+1^, written *c^i^* → *c*^*i*+1^, and (b) a transition from *c^i^* to any member of *T*(*c^i^*), written *c^i^* → *o ∊ T*(*c^i^*). From Bayes’ Theorem:

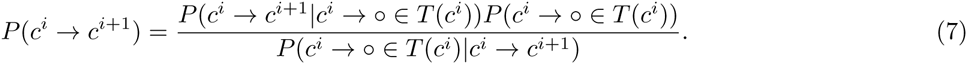

If *c*^*i*+1^ ∊ *T*(*c^i^*), the denominator *P*(*c^i^* → *o ∊ T*(*c^i^*) | *c^i^* → *c*^*i*+1^) = 1 and so

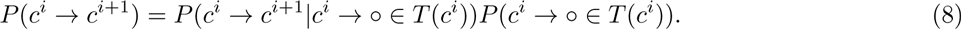

**Figure S1:**
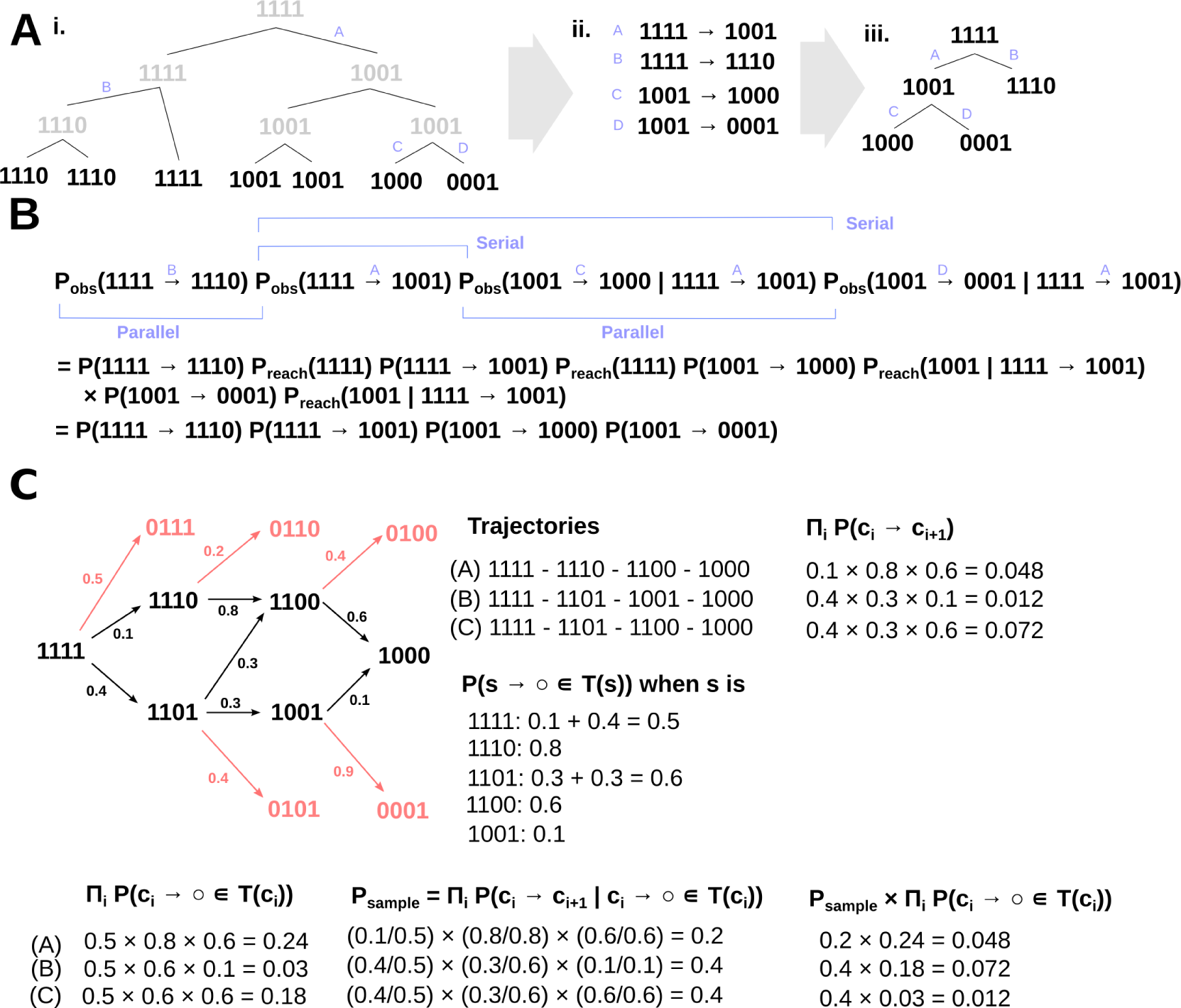
Construction of source data and structure of likelihood calculations in HyperTraPS – related to Fig. 1. (A) (i) A set of observed strings (black) and taxonomic relationships (lines) is used to infer strings throughout a taxonomic tree (grey). This taxonomy then reveals changes in strings between ancestor and descendant (blue letters, (ii)). This set of changes forms an evolutionary tree (iii) describing the mitochondrial evolution occurring ‘beneath’ the speciation and adaptation of ‘host’ organisms, which may involve branching lineages of different structure from the species taxonomy. (B) The likelihood of this tree under a model of branching random walkers is then computed. For clarity we use the symbol *P*(*a* → *b*) ≡ *P*(*b*|*a*; π). The product of each observation probability is written down, including terms due to serial descent down a single lineage and parallel branching. The dependence of observation probability on encountering the correct initial state drops out, as the lineage structure means that every observed ancestral state is the result of another observed transition. A product of transition probabilities *P*(*a* → *b*) ≡ *P*(*b*|*a*; π) then constitutes the likelihood function. Note that some transitions may involve changing more than one property (for example, A, 1111 → 1001), and may thus be accomplished through more than one trajectory. (C) HyperTraPS marginalisation and probabilities. An example transition network with edge weights is illustrated. Here, *s* = 1111 and *t* = 1000. All strings with a 0 in the first position are thus not *t*-compatible. These states, and transitions to them, are coloured pink. *t*-compatible states and steps are marked in black. No other transitions are supported. Three trajectories A, B, C lead from *s* to *t*. Their individual probabilities ∏_*i*_ *P*(*c_i_* → *c*_*i*+1_) are given at the top right. The other quantities involved in the main text are displayed for each trajectory, illustrating that the product of the sample probability of a trajectory *P_sample_*(*c*) and the product of the probability of making *t*-compatible steps at each state ∏_*i*_ *P*(*c^i^* → *o ∊ T*(*c^i^*)) exactly matches the trajectory’s individual probability.

If *c*^*i*+1^ ∉ *T*(*c^i^*), Eqn. 7 takes the form 0/0. However, this case involves a trajectory leading to a state that is not *t*-compatible, and thus cannot lead to *t*. Such a trajectory will therefore not contribute to the expression for *P*(*s* → *t*), which manifests mathematically as the observation that all subsequent steps *j* in such a trajectory will have *P*(*c^j^* → *o ∊ T*(*c^j^*)) = 0. Hence the product of probabilities associated with this trajectory interpreted as a path between *s* and *t* is zero.

By Lemma 1, we have that a trajectory leading from *s* to *t* must have *c*^*i*+1^ ∊ *T*(*c^i^*) for all *i*. In this case, the associated probability is the product of Eqn. 8 for each step in the trajectory:

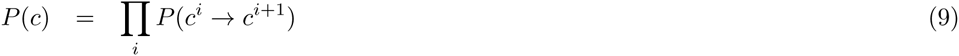

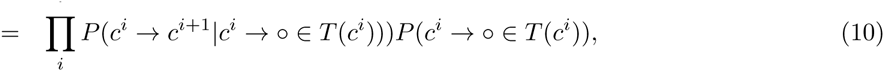

so the overall quantity of interest can be written

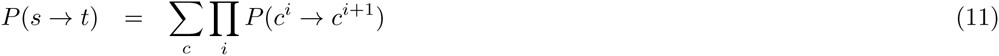

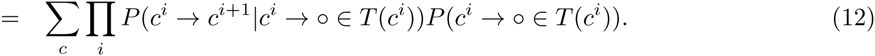

The idea behind this recasting is to facilitate a sampling approach. Consider simulating a trajectory, starting at *s* and progressing according to the following rule. At each state *c^i^*, identify the set of states *T*(*c^i^*) that may be reached by one acquisition that are *t*-compatible. Compute *p^j^* = *P*(*c^i^* → *c^j^*)/ ∑_*r∊T*(*c^i^*)_ *P*(*c^i^* → *r*) for each member *c^j^* of this set, where the sum over *r* is taken over all members of the set. Choose the next step according to the probabilities *p^j^*. This rule enforces t-compatibility in each step of the trajectory, thus forcing every trajectory to transition between *s* and *t*, while retaining the correct relative weighting of steps between states. *p^j^*, the probability of transitioning to state *c^j^* given that a transition is made to a *t*-compatible state, is identically *P*(*c^i^* → *c*^*i*+1^|*c^i^* → *o ∊ T*(*c^i^*)) when *j* = *i* +1. The factor of ∏_*i*_ *P*(*c^i^* → *c*^*i*+1^|*c^i^* → *o ∊ T*(*c^i^*)) is thus exactly the probability with which trajectory c will appear when simulations are performed over the set of trajectories constrained to begin at *s* and end at *t*. We can thus write

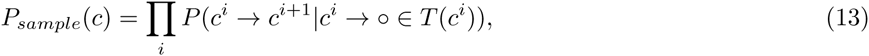

where *P_sample_(c)* is understood to mean the probability of simulating trajectory *c* given the above protocol. If we then adopt this sampling scheme and record the average value of ∏_*i*_ *P*(*c^i^* → *o ∊ T*(*c^i^*)) for each sample, we will obtain an estimate of the sum in Eqn. 12:

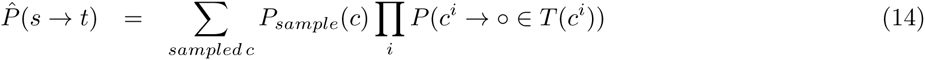

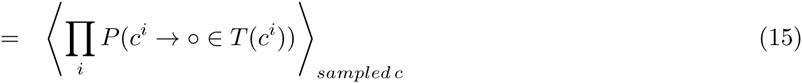

The advantage of this sampling approach is that we can simulate trajectories guaranteed to start at *s* and end at *t*, thus avoiding wasting computational time on trajectories that do not contribute to the overall sum. Convergence of 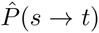 is therefore expected to proceed much more quickly than a naïve sampling approach involving unconstrained trajectories. An illustration of the quantities above for a specific transition network is shown in Fig. S1C.

### The HyperTraPS Algorithm

Here we describe our algorithm for hypercubic transition path sampling. We are not aware of this algorithm having been previously published; if this is true, we propose the name ‘HyperTraPS’, both standing for hypercubic transition path sampling and referring to the act of forcing trajectories towards specific points on a hypercube.

#### Algorithm 1 Hypercubic transition path sampling (HyperTraPS)

1. Initialise a set of *N_h_* trajectories at *s*.
2. For each trajectory *i* in the set of *N_h_*:

(a) Compute the probability of making a move to a *t*-compatible next step (for the first step, all trajectories are at the same point and the probability for each is thus the same); record this probability as 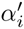.
(b) If current state is *s*, set 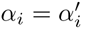, otherwise set 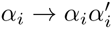
(c) Select one of the available *t*-compatible steps according to their relative weight. Update trajectory *i* by making this move.
3. End for each.
4. If current state (in all trajectories) is *t* go to 5, otherwise go to 2.
5. 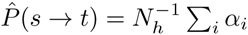.

In this algorithm, *α* is a vector, with each of *N_h_* elements progressively recording the product of probabilities П_*i*_*P*(*c^i^* → o ∈ *T*(*c^i^*)) of transitioning to any *t*-compatible state, where the product is taken over all steps so far performed in the corresponding trajectory. Each trajectory is simulated as above, by choosing a *t*-compatible transition at each state according to its relative weight. When *t* has been reached, the average П_*i*_*P*(*c^i^* → o ∈ *T*(*c^i^*)) is computed over all sampled trajectories.

*N_h_*, the number of sampled trajectories, is a parameter of the algorithm. Lower numbers will be computationally cheaper but will give a poorer sampling of possible trajectories and thus a less accurate estimate 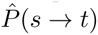.

### Inferring transition matrices

Experimental observations of mitochondrial genomes constitute a dataset 𝒟, which consists of a set of paired barcodes *a_k_, d_k_*, respectively the kth ancestral and descendant barcodes. We computed the likelihood 𝓛(π|𝒟) associated with a trial transition matrix π by sequentually using each *a_k_, d_k_* pair as the *s,t* pair in the HyperTraPS algorithm, and multiplying the likelihoods associated with each pair. For computational convenience, log-likelihoods *l* = log 𝓛 were used.

These log-likelihoods were then used in a Bayesian MCMC framework, with a new trial transition matrix π’ being produced from a current transition matrix π by applying Gaussian-distributed perturbations with standard deviation *σ* = 0.25 to each element of π. An initial parameterisation of π enforcing uniform gene loss probabilities with no evolutionary contingency was employed (in the *L* × (*L* + 1) representation, elements describing ‘base’ transition rates were all set to 1; all elements describing contingent modulation of these rates were set to 0). *N_h_* = 200 HyperTraPS trajectories were used to estimate likelihoods. 10^6^ MCMC iterations were used with a burn-in period of 2 x 10^5^ iterations, with posterior samples taken every 10^3^ iterations. *N_h_* and *σ* were chosen through preliminary investigation to lead to good chain mixing and convergence. *N_c_* = 10 repeats were performed with different random number seeds to check convergence and facilitate clean statistics. The posterior distributions on network parameters were visualised as described in the Main Text.

To reduce the search space of the inference process, we write π using π_*s→t*_ = *Z*^-1^ *P*(lose trait *i*|state *s*)*I*(*s; t; i*), where *I*(*s; t; i*) is an indicator function returning 1 if t is equivalent to s after the loss of trait *i*, and 0 otherwise, and *Z* is a normalising factor to ensure ∑_*t*_ π_*s→t*_ = 1. This structure allows us to coarse-grain the parameterisation of the problem by writing *P*(lose trait *i*|state *s*) = exp (*m*_*i*;1_ + ∑_*j m_i;j_* + 1_(1 – *s_j_*), where *m* is an *L* × (*L* + 1) matrix *m*, with elements in the first row *m*_*i*;1_ representing the ‘default’ probability of losing trait *i*, and elements in the subsequent rows *m_i;j_*_+1_ describing how this default probability changes in a state where trait *j* has already been lost. This representation, as discussed in Ref. [61] where this philosophy is also employed, allows us to use *O*(*L*^2^) parameters rather than the full *O*(2^*L*^) set of transition probabilities, while retaining the ability to model both independent gene loss propensities and potential contingencies of the loss of one gene on the presence of others. We illustrate its ability to satisfactorily capture evolutionary behaviour in the Supplementary Information.

## Validation

To test the performance of the HyperTraPS algorithm, we first considered the simplest possible case, where all possible transitions from every point are equally likely, where the start node *s* = 0^*L*^ is the node corresponding to an all-zero barcode, and where *t* is a single, specific target. The probability of encountering *t* is then the probability of sequentially acquiring the *n_t_* traits of *t* without acquiring any others. This probability is (*n_t_*/*L*) × ((*n_t_* – 1)/(*L* – 1)) ×…, hence:

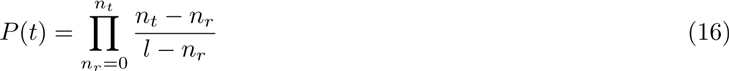

which is straightforwardly enumerable. We verified for a simple *L* =10 system that the HyperTraPS observation frequency exactly matched this analytic result for a range of random targets (Fig. S2A).

Next, we relaxed the restriction on π to consider more general transition matrices that lacked a straightforward analytic result for the observation probability of a given target. Once more, we considered the probability of observing a set of randomly chosen target strings, and compared the HyperTraPS results with those estimated from sampling many explicitly simulated random trajectories on the given hypercube. We fixed π at *L* = 10 and explored the correlation between the HyperTraPS and simulated results, which is excellent over many randomly chosen targets (Fig. S2B).

**Figure S2:**
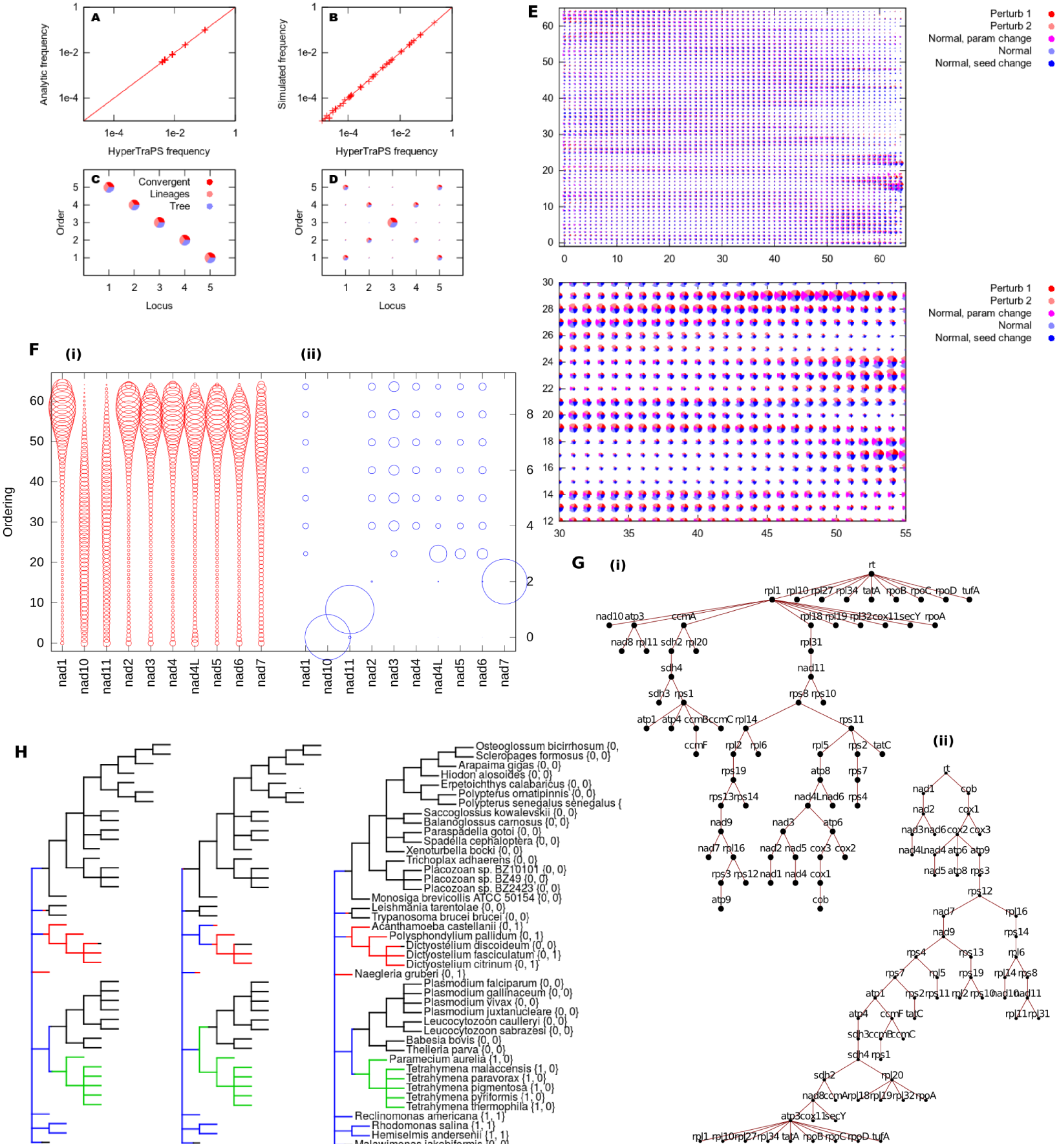
Validation of HyperTraPS algorithm and comparison to other approaches – related to Fig. 1. (A) Comparison of HyperTraPS observation frequency and analytic observation frequency for simple π and random single specific targets. (B) Comparison of HyperTraPS observation frequency and simulated observation frequency for non-trivial π and random targets. Departures at low frequencies are due to the finite sampling allowed by the simulation approach (2 x 10^6^ trajectories simulated for each target). (C-D) Reconstructed evolutionary dynamics using HyperTraPS and Bayesian inference, for (C) a simple underlying hypercube supporting a single evolutionary path and (D) a more complicated hypercube supporting two distinct pathways. A range of taxonomic relationships (covergent, linear, branching tree) between observations were considered in each case (see text). (E) Robustness of results with respect to perturbations in data gathering and simulation protocol. The posterior probabilities, represented by arc radii, associated with gene loss ordering for short illustrative simulations run with several different protocols. The ‘perturb’ label involves two randomly perturbed datasets as described in the text, testing robustness with respect to specific details of the phylogenetic and mtDNA structure reconstructions. The ‘normal’ involves the unperturbed dataset with default parameters; results using different simulation parameters (*N_h_* = 100, *σ* = 0.1) and a different random number seed are all shown. (top) Full 65 x 65 set of posteriors; (bottom) zoomed view on a particular region. Gene ordering (horizontal) is arbitrary in these plots and gene labels are omitted for clarity. All results are very comparable and do not lead to substantial changes in our general results. (F-H) Other related methods for trait evolution over related observations. (F) ‘OrderMutation’ inference of gene loss orderings given a reduced set of *nad[X]* genes and a reduced set of observations. (i) Shows the posterior distributions on loss orderings from our approach; (ii) shows the mean loss orderings from OrderMutation over 20 bootstrap resamples. (G) ‘Oncotrees’ inference of maximum likelihood trees of trait relationships. (i) Loss treated as the ‘acquisition’ of a loss event (root is 000…). (ii) Loss treated as explicit losses, inverting the normal action of oncotrees (root is 111…). In both cases, *cob, cox[X]* and many *nad[X]* genes are inferred to be lost late, and rare genes (for example, *secY*) are inferred to be lost early. Some structure corresponding to different complexes is visible: some *nad[X]* genes cluster together, as do some ribosomal genes. (H) ‘Simmap’ stochastic mapping of trait evolution on a reduced phylogeny. Three instances of stochastic maps of the evolution of { *nad10, nad11* } on a reduced phylogeny. Colours give states: {1,1} blue; {1, 0} green; {0,1} red; {0, 0, } black.

Finally, we tested the HyperTraPS algorithm in a controlled inferential setting. We constructed simple *L* = 5 artificial datasets consisting of samples from random walks on transition hypercubes with known transition probabilities, and attempted to infer these known underlying dynamics from the artificial data with HyperTraPS. The two transition hypercubes we used represented a simple case where a single evolutionary pathway is supported, and a more complicated case where two distinct pathways exist, and the trajectory experienced by a lineage is contigent on which of two equiprobable first steps is taken. For each transition hypercube, we constructed artificial data to cover a range of phylogenetic structures: totally convergent evolution (where no observed species is related); evolution involving the minimal number of possible lineages (one for a single pathway, two for two competing pathways); and a branching-tree phylogeny linking observed species in a more complicated arrangement. We thus covered a range of underlying evolutionary dynamics and a range of taxonomic connections between observations. In these cases, we used the reduced parameterisation of π described above (using *O*(*L*^2^) rather than *O*(2^*L*^) parameters while maintaining estimates of independent and contingent loss probabilities). Fig. S2C-D illustrate the excellent reconstruction of the dynamics supported by the underlying transition matrix in each case.

To confirm that our results for the mtDNA investigation were not dependent on specific details of the reconstructed phylogenetic tree from the NCBI Taxonomy Tool, or the inference of gene loss events, we applied random perturbations (changing presence/absence properties of individual genes with probability 0.05) to the set of ancestor-descendant pairs that arose from our data analysis. These random perturbations model the effects of changes to the phylogenetic structure and inferred ancestral mtDNA structures across our dataset. In parallel, to confirm that simulation runs were not becoming trapped and that chains were successfully mixing to converge on a true posterior, we compared the results from several different simulation protocols, involving different random number seeds and step sizes (see Fig. S2E). The agreement between the posteriors in all these cases confirms the convergence of the algorithm and its robustness to perturbations in the source data.

### Comparison with related approaches

The study of the evolution of traits across phylogenetically related lineages has an extensive history and associated literature (reviewed in Ref. [45]). Methods have been developed in this field to infer phylogenetic trees, to infer the evolutionary dynamics of traits on phylogenies, and to jointly infer both (a variety of each type are included in Ref. [71]). Often a single (continuous or discrete) trait or pairs of traits are considered; one approach that is particularly notable in the context of this study employs reversible-jump MCMC to explore the potentially correlated evolution of two discrete traits on a phylogeny [72]. However, we are unaware of an approach in this field that can consider the evolution of a large (*L* = 65) set of potentially interacting traits on a phylogeny. These approaches do provide alternative and sophisticated approaches for characterising uncertainty in phylogenies. However, we employ our straightforward and robust approach (see Methods) due to the taxonomic range of our study (and corresponding difficulty in using e.g. particular features like sequence alignments for phylogenies), our interest in an ordering of trait changes rather than an absolute temporal dimension, and our focus on trait dynamics rather than phylogenetic structure *per se*, which mtDNA studies have previously addressed [32].

A rather disconnected branch of literature attempts to describe the (usually irreversible) acquisition of coupled binary traits with time, usually given a set of independent observations of this process. This field is largely motivated by the target of inferring the dynamics of cancer progression, with the binary traits under consideration often corresponding to the presence or absence of chromosomal aberrations. Refs. [46] and [48] review and classify approaches to this question, which often involve assumptions about the causal links between traits (for example, that the network of trait acquisition influence is tree-like). Our method allows these assumptions to be relaxed to the case where traits can influence acquisitions arbitrarily, and in concert allows posterior ordering to be derived, facilitating subsequent exploration of connected factors.

The method proposed in Ref. [61] employs a similar Markov chain philosophy (and indeed uses the same strategy for reducing parameter space as in our method). This approach can be adapted to mirror the same likelihood function as HyperTraPS, by consider each transition between our mtDNA states as a start and end observation of traits. However, the likelihood calculation proposed in that study has a complexity of *O*(*nL*2^*k*^), where n is the number of observations, *L* the number of traits, and *k* a number of trait differences that separate two observations. As, in this study, *k* is sometimes of the same order of *L*, and *L* = 65, this likelihood calculation is intractable. Another approach, ‘phenotypic landscape inference’ [47], uses likelihoods estimated in time *O*(*nL*2^*r*^) (where *r* is the number of paths used to characterise a given transition), using an unbiased search which necessitated *r* ∼ *O*(2^*L*^) for satisfactory convergence (and many likelihood calculations to satisfactorily explore the parameter space of size 2^*L*^), and so is also intractable for *L* = 65. By comparison, the efficient probability-weighted algorithm in HyperTraPS has a complexity of *O*(*nk*2^*r*^), and the resultant polynomial rather than exponential scaling with *k* means that HyperTraPS provides a tractable (approximate) likelihood where the approaches of Refs. [61] and [47] would be intractable, making the investigation of these larger questions computationally feasible for the first time. Polynomial rather than exponential scaling in the number of traits means that progression dynamics can now be inferred for problems involving many traits without necessitating assumptions of tree or other structures underlying trait relationships (although such assumptions can straightforwardly be included by applying restrictive priors to the appropriate transition intensities). Additionally, as we demonstrate, increased computational speed facilitates fully Bayesian analyses of the dynamics of evolutionary/progression systems, allowing subsequent analysis of explanatory features and mechanisms, probabilistic statements about expected progression pathways from a given state, predictions about unmeasured trait values, and other important deliverables.

Fig. SS2F-H illustrates several existing approaches applied to our data on mtDNA gene loss. First, the ‘Oncotrees’ package (Fig. S2F), which attempts to infer a tree of relations between different traits that are acquired with time [73]. The trees that emerge from this approach are heuristically comparable to our findings: *cob* is notable at one limit of loss dynamics, with *cox[X]*, nad[X], and some *atp[X]* genes occupying points late in the ordering hierarchy, whereas rare genes like *sec Y* and *tat A* are present far lower. This picture provides independent confirmation for our findings, but does not characterise explicit posterior probabilities in the detail that our HyperTraPS-led approach does, and would not admit further analysis within the same framework to determine features that predict loss propensity. Second, the ‘OrderMutation’ approach of Ref. [74] (Fig. S2G), which attempts to infer the probability with which a given trait changes at a given ordering. This approach produces results comparable to ours when applied to an *n* = 10 subset of the genes we consider, but struggles with a larger *n* = 20 set and fails to process larger datasets. Third, the ‘Simmap’ approach [75, 71] (Fig. S2H), which characterises stochastic transitions between different trait patterns on a given phylogeny. This approach (when restricted to consider gene loss as irreversible) generates plausible realisations of evolutionary trajectories for small subsets of genes on our phylogeny (as illustrated in Fig. SS2F-H) and can characterise uncertainty in this trajectories. However, the 2^*L*^ different potential trait states in our mtDNA system, most of which are not observed in contemporary samples, rendered this approach intractable.

For ‘Oncotrees’, the full set of 2015 observations was treated as input. For ‘OrderMutation’, we use only the set of *n* = 74 unique genomes as input, and a reduced set of *L* = 10 genes from this set, namely {*nad1, nad10, nad11, nad2, nad3, nad4, nad4L, nad5, nad6, nad7*}, and bootstrap over 20 resamples, each using 10 simulations, to estimate mean orderings. For ‘Simmap’, a reduced phylogeny consisting of a subset of species was used for clarity (and to better illustrate diverse evolutionary trajectories, highlighting some unicellular branches); all branch lengths were set to 1. The genes {*nad10, nad11*} were chosen to reect diverse behaviour in this reduced phylogeny. Transitions that involved the acquisition of genes were prohibited by using a lower-diagonal transition matrix model. The output of ‘Simmap’ assigned equal rates (0.20) to each remaining possible transition between states, implying no asymmetry in transitions for this (reduced) example.

### Gene data acquisition and curation

Table S1 summarises the genetic and physical properties of genes that we consider in the model selection process.

### Compilation

The **length**\* (in bases) and **GC content** (average number of G bases and C bases per codon) of genes are taken straightforwardly from a sequence. The **GC skew**\* was computed as (*G* – *C*)/(*G* + *C*), where *G* is the number of G bases and *C* the number of C bases [12], from the gene sequence. The **strand\*** upon which a given gene was encoded was represented by a 0 (light strand) or 1 (heavy strand) according to GOBASE’s entry.

**Table S1.**
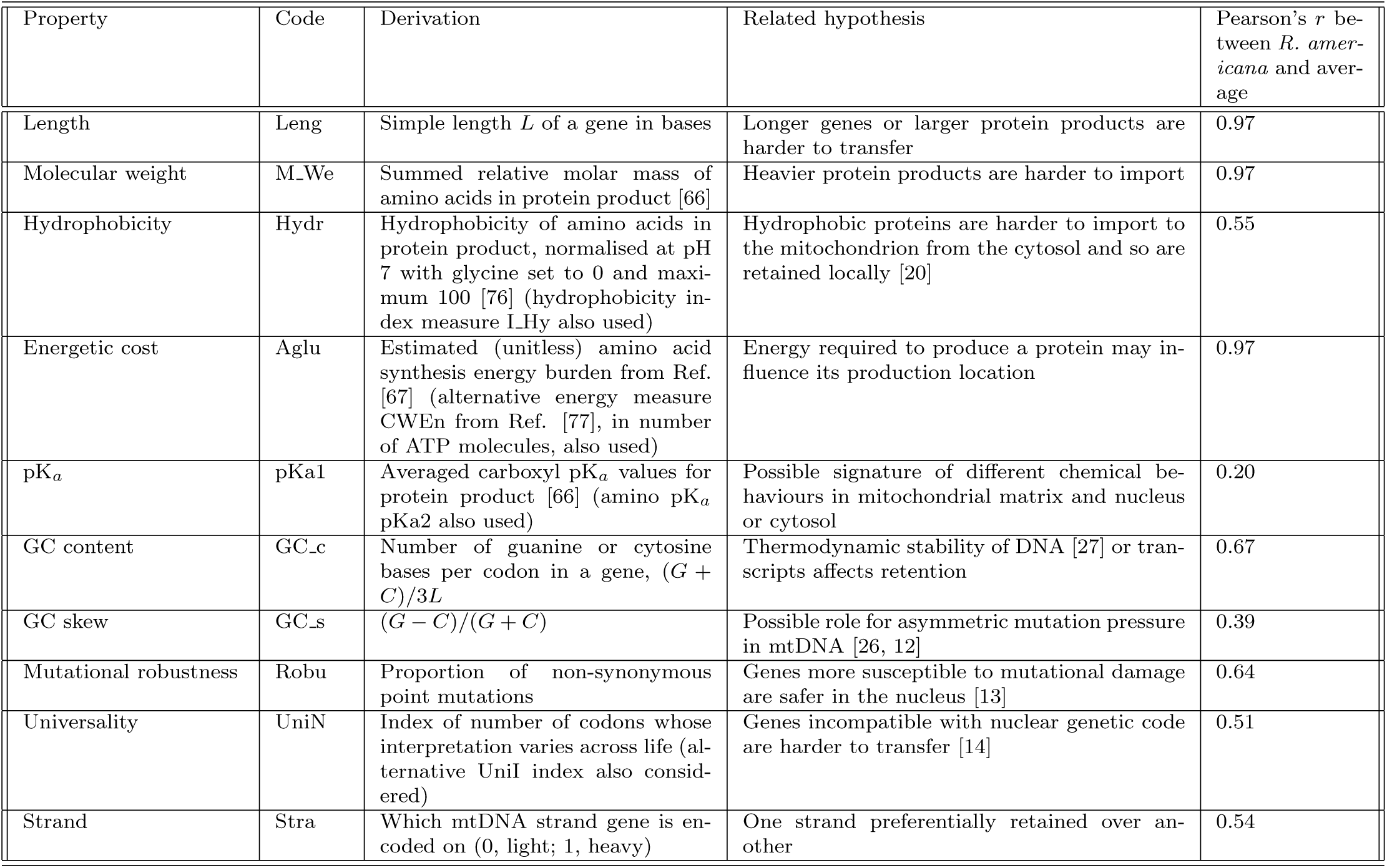
Gene properties and hypotheses compared in Bayesian model selection – related to Fig. 5. The different properties of genes and gene products used to explore hypotheses regarding mitochondrial gene retention. ETC, electron transport chain. Illustrative Pearson’s *r* correlation coefficient computed between the values of genes in *R. americana* and the values of genes averaged across all available eukaryotic genomes.

**Table S2.**
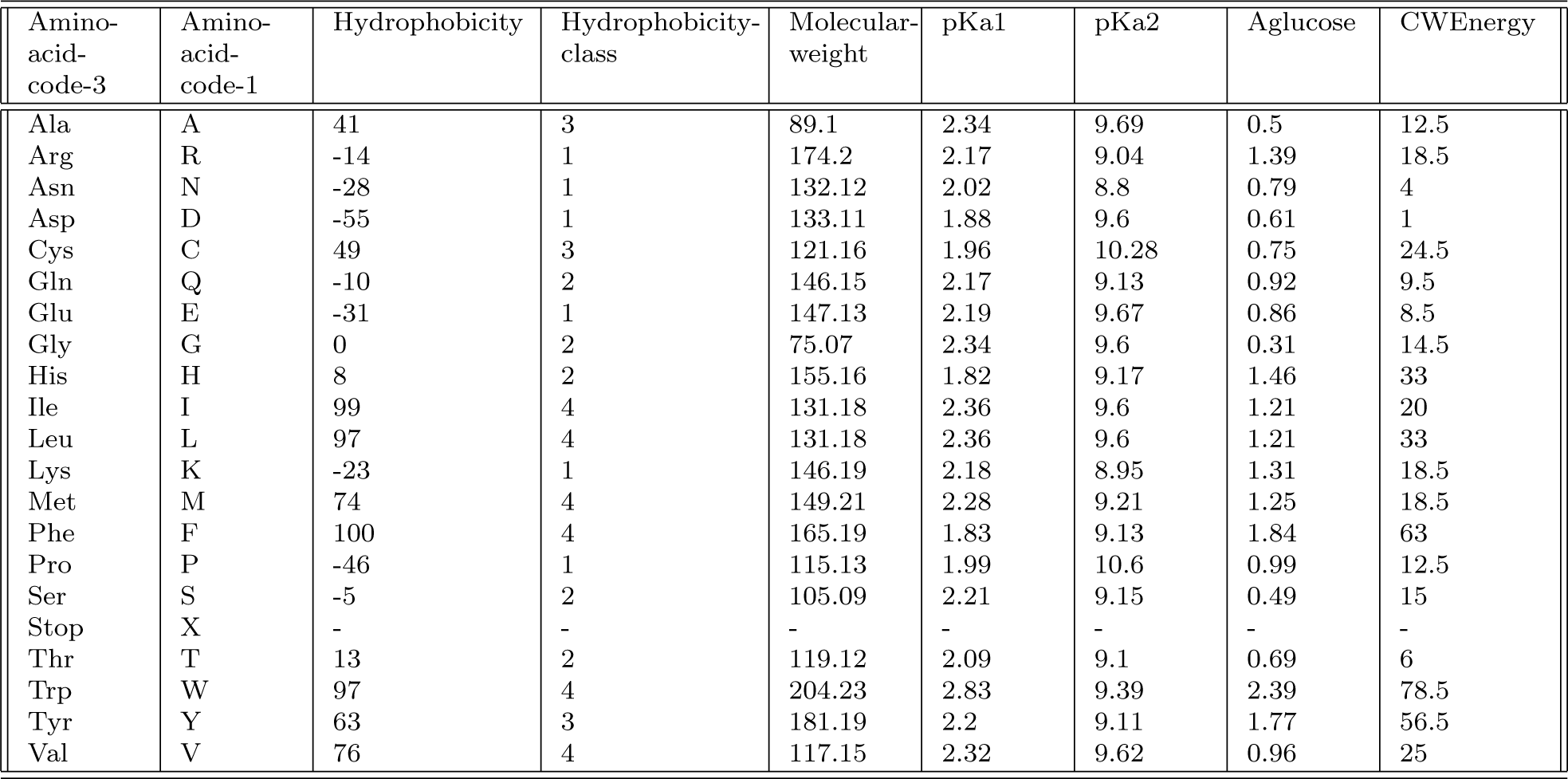
Amino acid properties used in model selection – related to Fig. 5. Numerical values of the properties described in the text. See text for sources.

**Table S3.** Non-universal codons – related to Fig. 5. The set of codons for which mitochondrial interpretation has been observed to differ in different taxa. The non-universality index counts the number of taxonomic cases for which such departure has been observed.

| Codon | Amino-acid-code-3 | Amino-acid-code-1 | Non-universal | Non-universal-index |
| --- | --- | --- | --- | --- |
| CUA | Leu | L | 1 | 1 |
| CUC | Leu | L | 1 | 1 |
| CUG | Leu | L | 1 | 1 |
| CUU | Leu | L | 1 | 1 |
| UAA | Stop | X | 1 | 1 |
| UCA | Ser | S | 1 | 1 |
| UAG | Stop | X | - (1) | - (2) |
| AAA | Lys | K | 1 | 3 |
| AUA | Ile | I | 1 | 5 |
| AGA | Arg | R | 1 | 7 |
| AGG | Arg | R | 1 | 7 |
| UGA | Trp | W | 1 | 9 |

Chemical properties of amino acids were taken from the compilation at http://www.sigmaaldrich.com/life-science/metabolomics/learning-center/amino-acid-reference-chart.html. The hydrophobicity and **hydrophobicity index** of a gene product was computed using this compilation (original data from Ref. [76]). **Amine group pK_a_**, carboxyl group pKa, and molecular weight* values were calculated using this compilation (original data from [62]).

**Glucose energy costs**\* were computed using the *A_glucose_* metric, based on the absolute nutrient cost required for amino acid biosynthesis, from Ref. [63]. Craig-Weber energy costs*, estimating the number of high-energy phosphate bonds and reducing hydrogen atoms required from the cellular energy pool to produce an amino acid, were taken from Ref. [77]. These biochemical properties are summarised in Table S2.

**Robustness** was assigned by counting the number of point mutations that would lead to nonsynonymous changes in the gene product. The **universality** and **universality index** of a gene were defined by considering the number of codons in the gene that were subject to different interpretations throughout the Tree of Life. These codons were identified using the NCBI compilation at http://www.ncbi.nlm.nih.gov/Taxonomy/Utils/wprintgc.cgi. The non-universality of a codon was set to 1 if its interpretation was ever different from the universal genetic code, and 0 otherwise; the non-universality index of a codon was equal to the number of different taxa in which it has been observed to differ (see Table S3).

Asterisks denote properties that are *not* averaged over gene length; it was deemed more appropriate to average other properties over genome length to gain a representative measure. To check for artefacts from this interpretation, we performed a (much more computationally demanding) model selection process including both the normalised and un-normalised values for each property; although coverage of individual models was unavoidably low in this procedure, the same consistent observation of GC content and hydrophobicity as important features was observed throughout.

The **assembly energy** of genes was quantified using the PDBePISA tool [66, 67] (http://www.ebi.ac.uk/pdbe/pisa/) to analyse the energetic interactions of the subunits in solved structures of electron transport chain complexes in the PDB [68] (specifically, Complex II (PDB 2h88, *Gallus gallus*), Complex III (PDB 3cxh, *Saccharomyces cerevisiae*), and Complex IV (PDB 1oco, *Bos taurus*)). The chains corresponding to products of genes in our reference set were identified, and for each chain, the total energy of all interfaces with other chains was recorded as its assembly energy.

We also explored the link between gene expression levels and evolutionary history of mitochondrial genes. The level of gene expression in animals has been postulated to affect the rate of sequence evolution of mitochondrial genes [28], possibly due to the deleterious effects of misfolding abundant proteins. To explore this link we required gene expression data of mtDNA genes, preferably in species with large numbers of mtDNA genes. We obtained RNA-seq data from *Phoenix dactylifera L.* (date palm) [65] and *Lolium perenne L.* (perennial ryegrass) [64], both containing ∼ 30 genes. We use data from mitochondria in *Lolium* and data from green leaves in *Phoenix*; other tissue types in the latter did not qualitatively change our findings.

Fig. S3D shows the links between gene expression levels in these species and inferred gene retention ordering and GC content in *R. americana* and across species. In *Lolium*, little correlation is present between gene expression and retention, or between expression and GC content. In *Phoenix*, we observe a moderate correlation between expression levels and gene retention. Stronger correlations are present between expression levels and GC content, particularly GC content from *R. americana.* This overall pattern of links is not compatible with a universal link between gene expression and mtDNA gene retention. However, it is compatible with a picture where a link between expression levels and gene retention is observed in some species where expression correlates with GC content (*Phoenix*), but not in other species (*Lolium*).

### Correlations and consistency

All properties except assembly energy were computed both using *R. americana* as a reference genome and by taking the average value across all organisms in which the given gene was present. Most properties (with GC skew a notable exception) showed at least reasonable correlations between these two approaches (Table S1), suggesting that these very coarse-grained mitochondrial genetic properties do not vary beyond recognition between organisms. GC skew displayed dramatic differences between different organisms, suggesting that other evolutionary pressures may act on this more detailed genetic feature [26, 12].

In Fig. S3D-E we illustrate the correlations between the different features used in this study. The strong correlations between the following physically similar pairs of features motivated the removal of one of the pair from the set of features explored in the main text: Hydr and I_Hy; pKa1 and pKa2; Aglu and CWEn; UniI and UniN.

We particularly focus on the connections between those features that we identify as key influences on mitochondrial gene loss: GC content, hydrophobicity, and energetic centrality. As discussed in the Main Text, these three features may intuitively be thought of as biologically connected: GC-rich codons encode hydrophobic amino acids, and hydrophobic peptide chains occupy central positions in complexes and within membranes. The weak and inverse (*r* = –0.34) correlation between GC content and hydrophobicity in the genes we consider (Fig. S3F) immediately suggests that this link may not represent the full story. Fig. S3F pursues possible links between these variables further, showing that none of these features has substantial predictive power for the others over the set of genes that we consider.

## Model selection

The values of gene properties across the set of *L* genes under consideration were collected and normalised to form a matrix *g_ij_*, where *g_ij_* is the normalised value of property *j* for gene *i*. This normalisation was performed by simply dividing each value by the maximum observed value for that property over all genes, thus ensuring that *g_ij_* ∈ [0,1].

Model selection proceeded by using a trial vector a to represent the coefficients of each of *N* features under consideration in a trial model. The sum

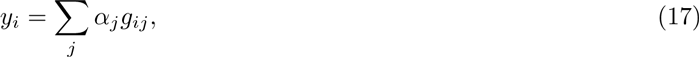

where *g_ij_* is the value of property *j* for gene *i*, was computed. The *L* genes under consideration were then ordered by ascending values of *y_i_*, yielding a vector of ranks *ρ*, where *ρ_i_* is the rank of gene i in the ordered list. Given the inferred posterior ordering *P_ij_* derived from HyperTraPS, we compute the likelihood associated with model *α* as 𝓛 = П_*i*_ *P_i ρi_*, the joint probability of each gene being lost in the order predicted by the model.

Bayesian MCMC was used to perform model selection given this likelihood definition. Two different prior protocols were used.

### Uniform priors

Each step, a uniform random number between 0 and 2^*N*^ was chosen. The binary representation of this number determined the features to be included in the model: if the ith bit in the binary representation of the number was zero, the ith element of a was set to zero. This procedure thus assigned a uniform prior distribution over each of 2^*N*^ possible model structures.

### Exponential priors

An exponentially-distributed random number with mean 2 was produced. This number gave the number of non-zero features to include in a trial model. This mean was chosen to allow a reasonable probability of choosing a full set of features, while also strongly favouring more parsimonious models.

Non-zero elements of a were assigned uniformly distributed random values on [—λ, λ], where λ = 5 x 10^3^ was chosen to exceed the range of model parameterisations found through a preliminary bootstrapping investigation. MCMC was used to sample from the posterior distribution of model structures and model parameterisations. 10^9^ samples were performed, chosen to give sufficient coverage to each of the 2^13^ ⋍ 10^5^ possible models. Figs. 3A-C demonstrate the consistent favouring of GC content and hydrophobicity across this set of protocols (different priors and different sizes of feature set).

The Main Text shows the correlation between inferred gene loss ordering and the predictions of a statistically-supported model involving GC content and hydrophobicity. Fig. S3G demonstrates that these predictions also correlate well with direct biological observations – specifically, the number of distinct mtDNA structures that contain a given gene. Thus, genes with high GC content and high hydrophobicity are observed in many different mtDNA structures; those with low GC and hydrophobicity are observed much less frequently.

In reporting a p-value associated with the link between a given model and inferred retention properties, the fact that we have selected one of many models and used a fit parameterisation must be taken into account. However, even given the substantial multiple hypothesis correction required when dealing with 2^10^ different model structures, the low p-value of 2 x 10^−11^ provides a compelling suggestion that the link between GC content, hydrophobicity, and retention is not coincidental.

**Figure S3:**
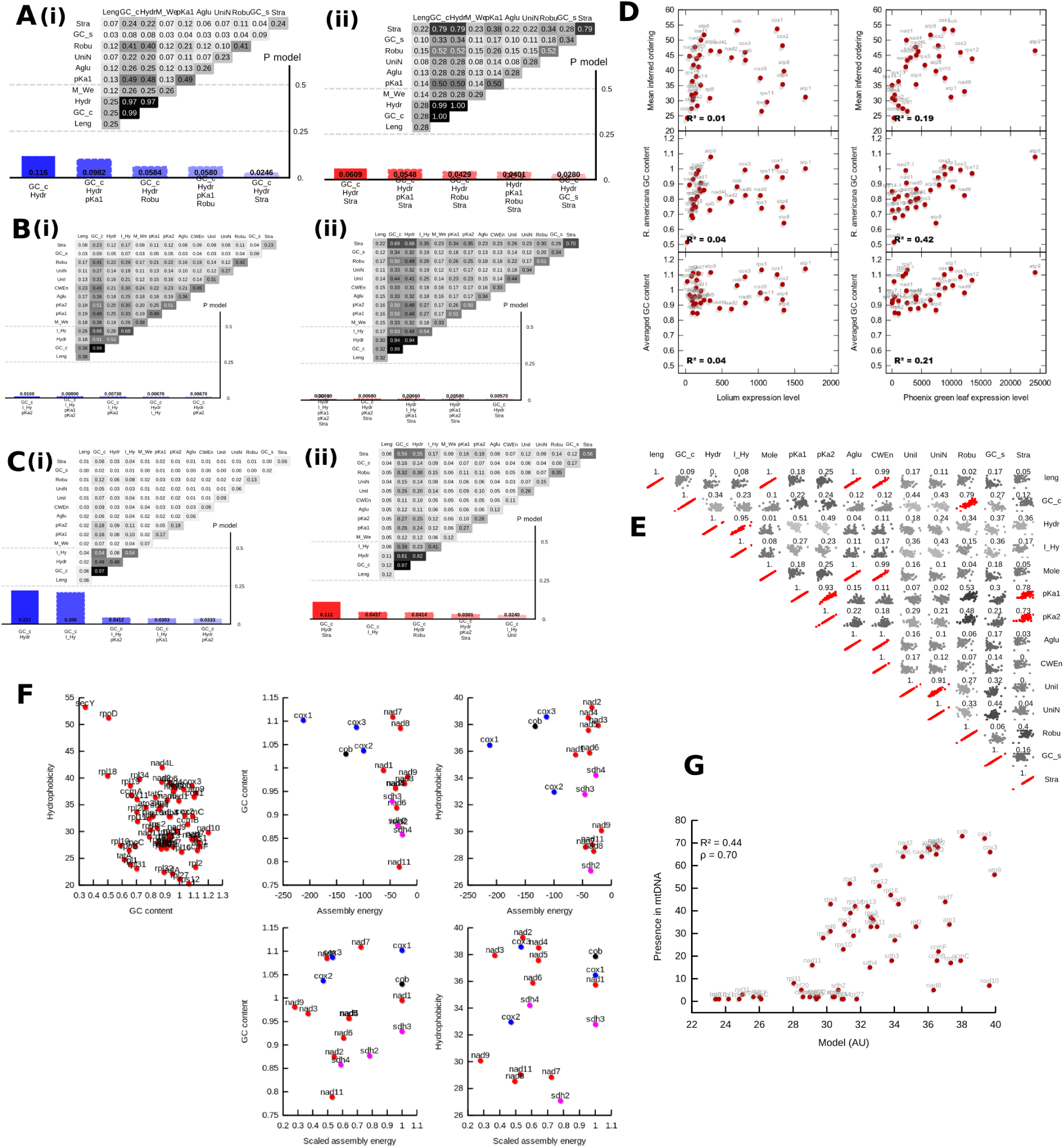
Model selection under different protocols, and correlations and links between gene features – related to Fig. 5. (A) Uniform priors on model structure with default feature set; (B) uniform priors on model structure with expanded feature set; (C) exponential priors on model structure with expanded feature set. As before, columns show the support for the most supported model structures, and matrices show the proportion of inferred models in which a given feature (diagonal elements) or pair of features (off-diagonal elements) occurs; (i) show model selection based on *R. americana* features, (ii) based on taxa-average features. (D) Correlations between gene expression levels, inferred loss ordering, and GC content in two plant species (see text). (E) Correlations between different model selection features. Scatter plots (axes omitted) and Pearson’s r for correlations between pairs of features used in model selection, using the values derived from cross-species averaging. (F) Correlations between features identified as linked to gene loss. Relationships between GC content, hydrophobicity, and assembly energy for all genes considered (GC vs hydrophobicity, as in the corresponding element of Fig. 8E) and ETC subunits (assembly energy). Scaled assembly energy gives each subunit’s energy scaled by the highest-magnitude energy within that protein complex. Little correlation exists between any variables; the *cox[X]* and *sdh[X]* genes display a moderate link between GC content and scaled assembly energy (bottom centre) but the null hypothesis of no correlation cannot be discarded (*p* > 0.05 in both cases). GC content and hydrophobicity are unitless; unscaled assembly energy has units of kJ mol^-1^. (G) Correlation between fitted model and mtDNA observation patterns. As in the Main Text, a model involving GC content and hydrophobicity scores for each gene was constructed and parameterised to fit the inferred mean of gene loss ordering. This plot shows values for each gene under that parameterised model, against the number of distinct mtDNA structures observed in our dataset that contain that gene. GC content and hydrophobicity thus exhibit a strong link to observed gene occurrences, as well as loss ordering (Main Text).

## MtDNA-wide GC content across taxa

As discussed in the text, our analysis provides strong support for genes with high GC content *relative to other genes in the same organism* being preferentially retained in mtDNA. Patterns of GC content vary dramatically between species [27]. Notably, at a sequence level, we have found that protein-coding genes in mtDNA can in some species display a bias *against* GC content, in the sense that GC-poor codons are used more often than GC-rich codons to encode a given amino acid (see below). However, the observation that GC-rich codons are less frequent in the mtDNA of some species does not necessarily conflict with our finding that genes with *relatively* high GC content are preferentially retained. Asymmetric mutational pressure generally acts at the sequence level to reduce GC content and enrich GC-poor codons in organellar genomes [26]. We propose that selective pressure at the structural level generally acts to retain genes with higher GC contents, against a background reduction in GC content across the mtDNA genome from asymmetric mutation. Our results thus suggest a tension between an entropic mutational drive at the sequence level (decreasing GC content in codons) and a selective drive at the genomic level (retaining genes with higher GC content).

To investigate trends in GC content throughout the mitochondrial genomes of different species, we first computed a ‘null model’ describing the expected pattern of codon usage if every codon encoding a given amino acid was used in the genome with equal probability (no favouring of one codon over another). We then recorded the actual patterns of codon usage in individual mtDNA genomes from sequence data, and compared the two. In Fig. 5D we illustrate the observation that *R. americana* displays an observable bias against those codons with high GC content, preferentially using codons with lower GC content to encode a given amino acid. No such bias exists in *H. sapiens.* This variability in GC codon bias is observed across taxa.

The observation of species displaying a genome-wide bias against high GC codons are not incompatible with our findings that genes with high GC content are preferentially retained in mtDNA. A genome-wide pressure against high GC codons can exist independently of an inter-gene favouring for high GC content. Thus, even if an organism’s environment or biochemistry strongly favours low GC codons throughout the full mtDNA sequence, our statement that genes with a *relatively* high GC content are retained is unaffected.

One reason for the observed link between GC content and retention could be an indirect relationship through different levels of structural conservation between mtDNA genes. In this picture, asymmetric mutation pressure drives a reduction in GC content, but structural constraints are stronger in highly conserved genes, meaning that these genes retain high GC content. If highly conserved genes are preferentially retained in mtDNA (perhaps due to their structural importance in complex assembly), highly retained genes will passively display higher GC content.

To test this hypothesis we examined the GC content at different positions within the (less synonymous) first and second positions and (more synonymous) third positions in each codon in mtDNA genes, reasoning that structural conservation on a background of asymmetric mutation pressure would lead to higher GC content at nonsynonymous sits. In *R. americana*, we do observe (Fig. 5C) a notably lower GC content in synonymous loci, suggesting that (assuming a uniform initial condition) nonsynonymous loci are under higher selective pressure to retain their GC content. However, this behaviour is less clear in taxa-averaged gene data and vanishes for *H. sapiens* (Fig. 5C), suggesting that across taxa the link between GC content and simple structural conservation is less pronounced than in *R. americana*.

We sought to determine how much a potential link between structural conservation and mtDNA gene retention could explain the observed signal associated with GC content. To explore this question, we included both total GC content and position-specific GC content in our model selection process. If gene retention propensity is solely determined by degree of structural conservation, we would expect GC content at nonsynonymous positions (interpreted as a proxy for conservation) to be favoured over total GC content in model selection. Instead, we see an even combination of total GC content and nonsynonymous GC content in *R. americana*, and a pronounced favouring of total GC content in taxa-averaged gene properties (Fig. S4A-B). We conclude that while a link between structural conservation and gene retention can explain some of the signal associated with GC content, it cannot be solely responsible for the appearance of this signal across species.

## Notes and extensions

### Signal emission probabilities

We have assumed throughout that the probability associated with emitting signals that correspond to observations provides only a multiplicative constant factor independent of a specific model parameterisation. Here we discuss and illustrate the details of these emission probabilities.

**Figure S4:**
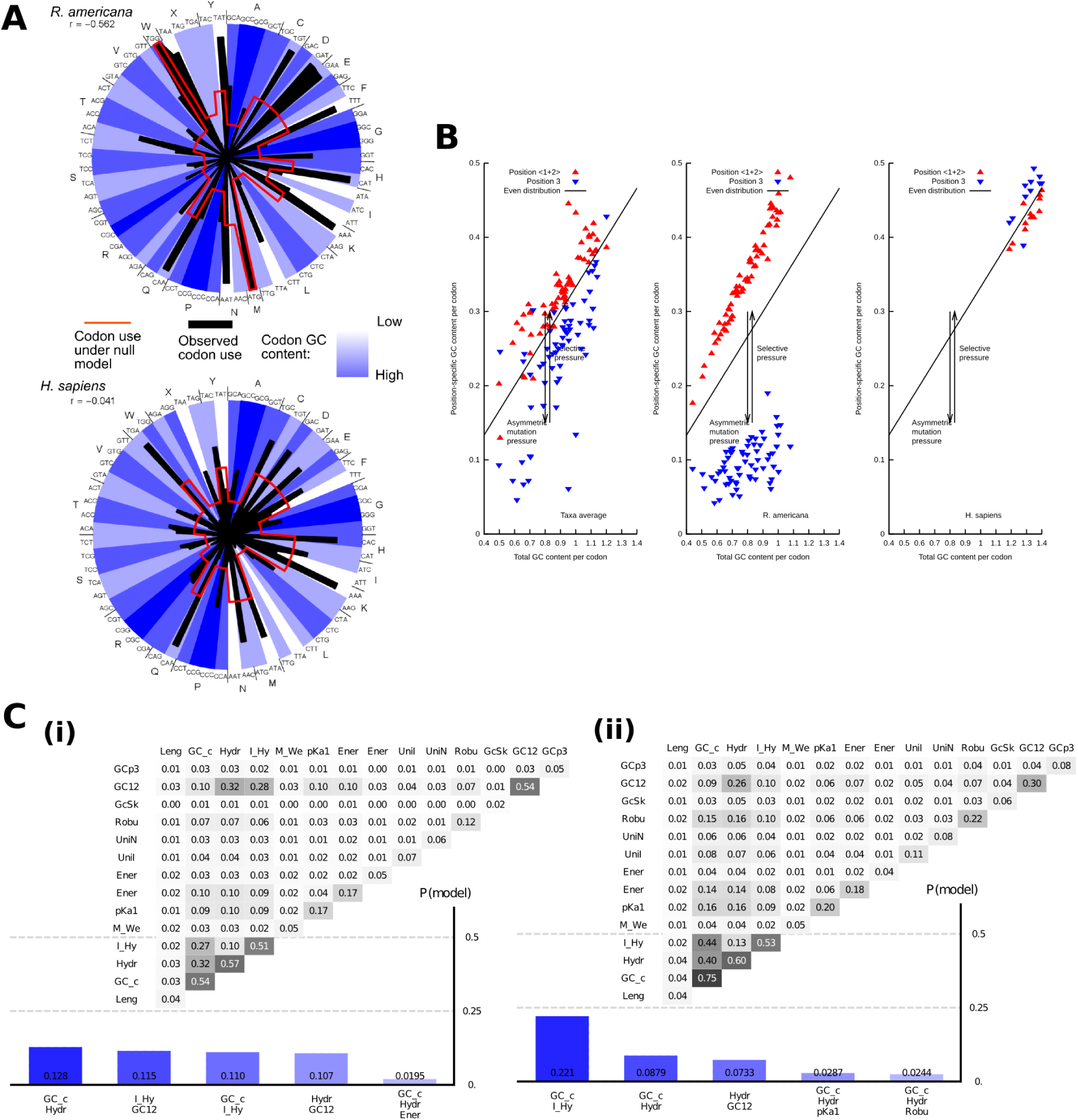
Model selection with different GC statistics – related to Fig. 5. Model selection using nonsynonymous (GC12) and synonymous (GCp3) GC content as well as total GC content (GC_c) in (A) *R. americana* and (B) taxa-averaged data. Total GC content still plats a dominant role in the most explanatory models.

In the first application of phenotype landscape inference [47], observations were independent (due to convergent evolution) and consisted of individual signals, sometimes containing uncertain data. Recall that in our mathematical model, signals are emitted uniformly from an evolving system with a given probability *P_em_ission* at each state. The probability of making a speciic observation is then the probability of the system being in a state compatible with the observation when a signal is emitted. As *P_emission_* was a constant of the model, this observation probability is straightforwardly proportional to the number of states in sampled trajectories that match that observation.

We now discuss the emission of signals down a lineage in more detail. Consider the case where we have π observations known to arise from the same lineage, and that these observations are time-ordered. What is the probability with which a system, randomly emitting signals, emits a set of signals at the speciic times matching an observation?

To compute this probability, we irst assume that the system will emit exactly π signals over a trajectory of length *L*, and that the emission of these signals is random and uniform over the trajectory. The probability of emitting exactly *n* states is a function of the assumed emission rate A alone, independent of the data and transition network. We write *P_λ_(n)* for this probability.

Assuming that π signals are emitted, and thus picking up a factor *P_λ_(n)* in our overall probability, we can represent the times of signal emission as *n* independent random variables taking values between 0 and *L* – 1, where each value corresponds to the number of steps from the origin state that have occurred when a signal is emitted. We are interested in the probability with which, when ordered, the set of these timings matches the observation pattern we require. We write this pattern as s = {*s*_1_,…,*s_n_*} where *s*_1_ ≤ *s*_2_ ≤… ≥ *s_n_*, with each element again corresponding to the number of steps from the origin state at which an observation is made.

Denote by *m* the number of distinct values taken by the elements of s, and by *c*_1_, *c*_2_, *c_m_* the number of times each of these values occurs. Thus, s = {0,0,1, 2, 3, 3} has *m* = 4 and c = {2,1,1, 2}. The number of emission patterns that lead to the observation of c upon time ordering is the number of ways that this set can be ordered, taking identical values into account, which is 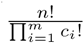. The probability of observing a given s is then this degeneracy multiplied by the probability with which each value is sampled:

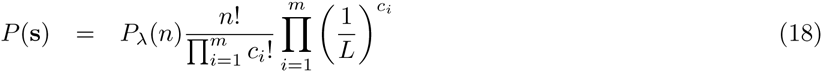

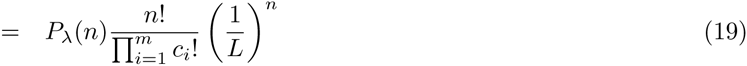

When each observation is known to have occurred at speciic states, the ordering is ixed, and hence *P*(s) is a constant. For example, the observations 111 → 101, 101 → 000 correspond to s = {0,1, 3}. We then have

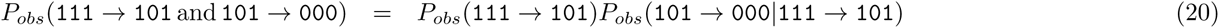

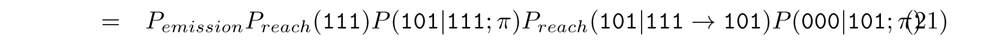

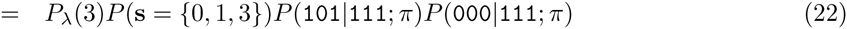

When an observation contains uncertain traits, the contributions from different observation patterns need to be accounted for. For example, the observation 11* can be matched by a signal with s = {0} emitting 111 and by a signal with s = {1} emitting 110. We then have

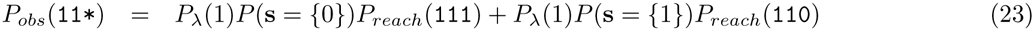

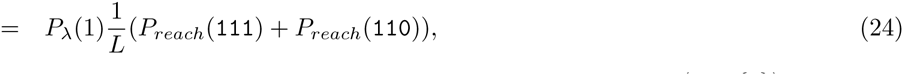

where the second line follows because, in the case of single observations for each lineage, *P*(s = {*c*}) is equal to 1/*L* for all *c*, as the probability of emission from any single state is uniform.

To illustrate the structure of probability calculations associated with emission probabilities, consider a system constrained to always follow the trajectory 111 → 011 → 001 → 000. The probabilities associated with several different observations, and the possible sets of states giving rise to each observation, are illustrated in Table S4.

### Incomplete data

We have discussed the case of complete data observed in parallel and/or serial descents pattern, and incomplete data observed in parallel descent patterns. A natural question is how to deal with incomplete data in serial descent patterns. We aim to address this more complicated question in further work; it is not required for the original application of this approach [46] (where evolution was completely parallel) or the application in this paper (where data is complete).

Several complications prevent the natural extension of the above analysis to lineages containing incomplete data. Firstly, ancestral properties inferred from incomplete contemporary measurements may themselves be incomplete. For example, if two descendents have properties *11 and 01*, the most reasonable inference of the properties of their ancestor is *11.

**Table S4.**
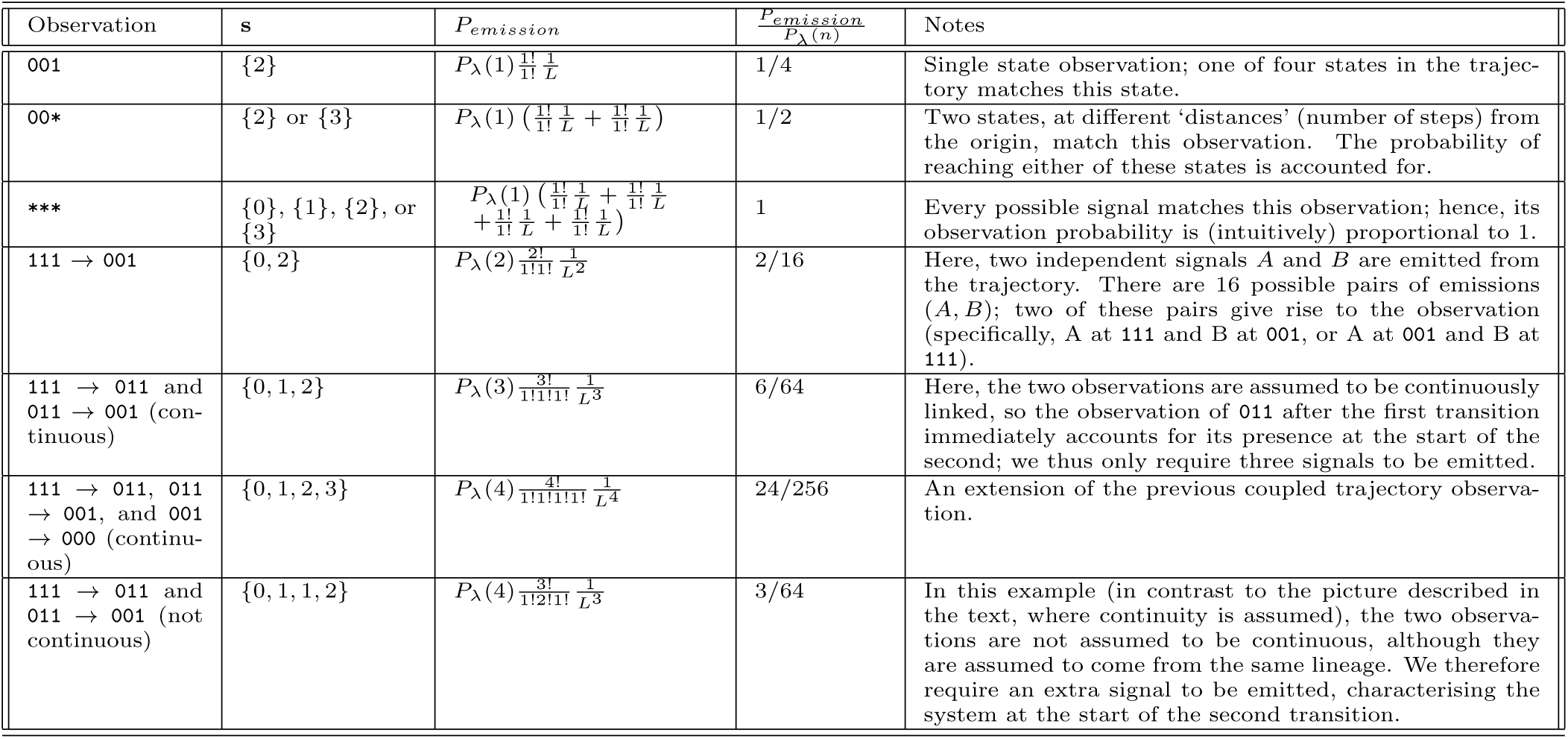
Probabilities of emission patterns for specific targets – related to Fig. 1. Here we demonstrate emission probabilities corresponding to different observations. We constrain evolution to follow the trajectory 111→011→001→000. This has the effect of removing transition probability factors from observation probabilities, as each transition has probability 1. The remaining observation probabilities thus depend only on the system emitting signals at appropriate points in the trajectory. For each observation, the pattern of signals s that give rise to that observation is shown, and the emission probability *P_emission_* from Eqn. 19 is given. We note that for more general transition networks, many more trajectories would be supported, and transition terms from each, not in general reducing to 1, would have to be included: accounting for these terms is the focus of the majority of the previous analysis.

Secondly, when computing observation probabilities involving an initial state with incomplete data, we cannot be certain that that initial state has been reached by a previous transition in the lineage under consideration. Consider a lineage in which a step is known to have identified a state in the set *U(t)*. We are interested in the probability of observing a transition to *b*; recall that *P_obs_*(*a* → *b*) = *P*(*b*|*a*; π)*P_reach_*(*a*; π). If *a* ∊ *U(t)*, *P_reach_*(*a*; π) will be nonzero but in general not unity. We therefore need to consider ∑_*a*_’_∊*U(t)*_ *P_obs_*(*a*’ → *b*) = *P*(*b*|*a*’; π)*P_reach_*(*a*’; π).

Thirdly, the patterns of signal emission are rather more complicated in the case of uncertain data. Previously, the structure of s, denoting the stages at which signals are emitted, was either ixed for each independent lineage (for example, Eqn. 22) or consisted of single elements (for example, Eqn. 23). For lineages involving several incomplete observations, we will in general have a large number of possible s structures, each involving several different elements. Furthermore, the time ordering of assumed signal emission times may vary according to the speciic set of states being considered as responsible for those signals. Extending the above calculations to account for all possible emission pattern options in these cases will be the central focus of future extensions of this work.

For now, we note that the simplest possible incarnation of this general approach will be equally applicable to cases with incomplete data. That is, simulating evolutionary trajectories and random signal emission from each, then comparing the emission of signals to observed data, will yield an estimate of the likelihood associated with a set of observations. However, the simplifying steps applied in the above analysis, such as efficient path location using HyperTraPS, the neglection of emission probabilities as constant multiplicative factors, and assumptions of continuous time-ordered progression through lineage observations, may cease to hold in this more complicated case, and the straightforward simulation approach may thus be computationally demanding. Further work will identify simplifying approaches and efficient simulation protocols to use in this context.

